# GWAS identifies candidate regulators of in planta regeneration in Populus trichocarpa

**DOI:** 10.1101/2022.06.08.495082

**Authors:** Michael F. Nagle, Jialin Yuan, Damanpreet Kaur, Cathleen Ma, Ekaterina Peremyslova, Yuan Jiang, Alexa Niño de Rivera, Sara Jawdy, Jin-Gui Chen, Kai Feng, Timothy B. Yates, Gerald A. Tuskan, Wellington Muchero, Li Fuxin, Steven H. Strauss

**Affiliations:** Department of Forest Ecosystems and Society, Oregon State University, Corvallis, OR, USA; Department of Electrical Engineering and Computer Science, Oregon State University, Corvallis, OR, USA; Statistics Department, Oregon State University, Corvallis, OR, USA; Biosciences Division, Oak Ridge National Laboratory, Oak Ridge, TN, USA; Center for Bioenergy Innovation, Oak Ridge National Laboratory, Oak Ridge, TN, USA; Bredesen Center for Interdisciplinary Research, University of Tennessee, Knoxville, TN, USA

## Abstract

Plant regeneration is an important dimension of plant propagation, and a key step in the production of transgenic plants. However, regeneration capacity varies widely among genotypes and species, the molecular basis of which is largely unknown. While association mapping methods such as genome-wide association studies (GWAS) have long demonstrated abilities to help uncover the genetic basis of trait variation in plants, the power of these methods relies on the accuracy and scale of phenotypic data used. To enable a largescale GWAS of *in planta* regeneration in model tree *Populus*, we implemented a workflow involving semantic segmentation to quantify regenerating plant tissues (callus and shoot) over time. We found the resulting statistics are of highly non-normal distributions, which necessitated transformations or permutations to avoid violating assumptions of linear models used in GWAS. While transformations can lead to a loss of statistical power, we demonstrate that this can be mitigated by the application of the Augmented Rank Truncation method, or avoided altogether using the Multi-Threaded Monte Carlo SNP-set (Sequence) Kernel Association Test to compute empirical *p*-values in GWAS. We report over 200 statistically supported candidate genes, with top candidates including regulators of cell adhesion, stress signaling, and hormone signaling pathways, as well as other diverse functions. We demonstrate that sensitive genetic discovery for complex developmental traits can be enabled by a workflow based on computer vision and adaptation of several statistical approaches necessitated by to the complexity of regeneration trait expression and distribution.

## Introduction

Plant genetic engineering and gene editing have produced new varieties of crops with a variety of valuable traits (NAS, 2016; Jaganathan *et al*., 2018). However, the ability to impart new traits by these methods is limited to crop species with genotypes that can reliably undergo transformation and regeneration (TR). TR requires developmental responses to a series of hormone treatments and amenability to gene insertion, and the capacity for both varies greatly between and within species (Altpeter *et al*., 2016). The causes of this great variation in recalcitrance are poorly known; however, GWAS – with its potential to identify genes whose variation plays a key role in capacity for TR – can greatly enhance understanding of the TR process. In addition, the identified genes may serve as “reagents” for overcoming recalcitrance, similar to how overexpression of morphogenic regulator (MR) genes can enhance *in vitro* regeneration of transgenic shoot in a variety of species (Gordon-Kamm *et al*., 2019). *In planta* transformation methods can also be enhanced by MR genes, including those in *Populus tomentosa* (Deng *et al*., 2009), *Nicotiana benthamiana*, tomato, potato and grape (Maher *et al*., 2020). However, given the complexity and genotypic variation in TR capacity, it is likely that only a fraction of the potentially useful MR genes have been identified.

To help identify the genes responsible for variation in TR, we conducted GWAS in a population of 1,219 wild cottonwoods that had been re-sequenced by the US Department of Energy, up to 917 of which were previously studied for a variety of traits (Zhang *et al*., 2018a; Tuskan *et al*., 2018; Muchero *et al*., 2018; Bdeir *et al*., 2019; Weighill *et al*., 2019; Chhetri *et al*., 2020). We focused on regeneration from cut stems, while considering these may be a direct substrate for accelerated *in planta* transformation systems, and because of the expectation that regeneration processes are likely to share many elements whether induced *in planta* or *in vitro*. GWAS and related association mapping methods have previously been applied to study variation in the rates of *in vitro* regeneration in Arabidopsis, rice, wheat, rose, sorghum and poplar, among other plants (reviewed by Lardon & Geelen, 2020).

Regeneration phenotypes are notoriously difficult to quantify, whether *in planta* or *in vitro*. Calli and emerging shoots are often highly variable and complex in shape, color, and size, and sequential measurements are hard to take without damaging or contaminating regenerating tissues. This appears to have limited sample sizes in prior GWAS studies of regeneration. For example, Tuskan et al. (2018) selected only 280 genotypes to phenotype callus growth from a re-sequenced GWAS population of 1,084 *P. trichocarpa* genotypes. A similar GWAS of callus differentiation into shoots in *P. euphratica* was limited to 297 genotypes (Zhang *et al*., 2020). Nguyen et al. (2020) noted the “extremely laborious” nature of phenotyping *in vitro* traits as a consideration in their GWAS of callus formation across 96 rose genotypes (Nguyen *et al*., 2020).

Because of the importance of a large and precise sample for statistical power in GWAS (López-Cortegano & Caballero, 2019), we developed a computer vision method to measure regeneration from sequential images of decapitated, regenerating stems. Over 40 published studies have made use of diverse computer vision methods in GWAS of plants, including Arabidopsis, maize, wheat, rice, sorghum, soybean and barley, among others (Guo *et al*., 2022; Affortit *et al*., 2022; reviewed by Xiao *et al*., 2021). Developmental traits related to biomass and growth of whole plants or specific tissues have commonly been studied using various methods for segmentation of RGB images, including those involving color thresholds (Yang *et al*., 2015; Das *et al*., 2015; Al-Tamimi *et al*., 2016; Guo *et al*., 2018; Gage *et al*., 2018; Pham *et al*., 2019; Seethepalli *et al*., 2020; Campbell *et al*., 2020; Ogawa *et al*., 2021; Affortit *et al*., 2022), machine learning approaches such as support vector machines and random forests (Zou *et al*., 2019; She *et al*., 2019; Carlier *et al*., 2022; Xu *et al*., 2022), and deep convolutional neural networks (DCNNs; Ma *et al*., 2019a; Yasrab *et al*., 2019; Xu *et al*., 2022; Kamath *et al*., 2022). In recent years, DCNNs similar to that employed in the present study have demonstrated abilities to outperform earlier methods (Adams *et al*., 2020) and have become the dominant approach used for diverse computer vision tasks. In the context of plant phenotyping, this was demonstrated by the unparalleled performance of neural networks for the Leaf Segmentation Challenge benchmark dataset (Aich & Stavness, 2017; Dobrescu *et al*., 2017).

We report a GWAS of *in planta* regeneration in poplar, using a computer vision workflow based on deep segmentation to quantify specific regenerating tissues, and four association mapping methods to identify genetic markers associated with regeneration traits. The non-normal and skewed characteristics of segmentation-based traits presented a challenge for GWAS, which commonly relies on an assumption of normally-distributed residuals. Due to this, we tested multiple GWAS methods with adaptations to avoid violating this assumption while also preserving statistical power. We found over 200 statistically-supported gene candidates with diverse physiological roles that include hormone signaling, plant stress response, control of cell division, and cell wall structure – as well as many genes of unknown function.

## Materials and Methods

### Experimental design

The primary objective of our work here was to develop, test, and refine workflows using computer vision to measure developing tissues during regeneration. The system also needed to be sensitive enough to enable detailed genetic characterization of regeneration traits. Toward this end, we first adapted a DCNN of the PSPNet architecture to compute segmentation masks for regenerating plant tissues (callus, shoot and unregenerated stem) on a high-throughput scale. Second, we evaluated the statistical considerations of using these traits in GWAS, whether by models often relying on trait transformation to avoid violating assumptions of normally-distributed residuals, or by models that avoid this assumption altogether by binarization or permutation and computation of empirical *p*- values. Finally, we applied these methods in a population with over 1,200 wild poplar genotypes ideal for GWAS due to a remarkable number and density of single-nucleotide polymorphisms (SNPs; up 34 million depending on GWAS method) and extremely low linkage disequilibrium (LD; Zhang *et al*., 2018a; Tuskan *et al*., 2018; Muchero *et al*., 2018; Bdeir *et al*., 2019; Weighill *et al*., 2019; Chhetri *et al*., 2020). An overview of the experimental population and analysis pipeline is shown in Fig. 1.

**Figure 1.**
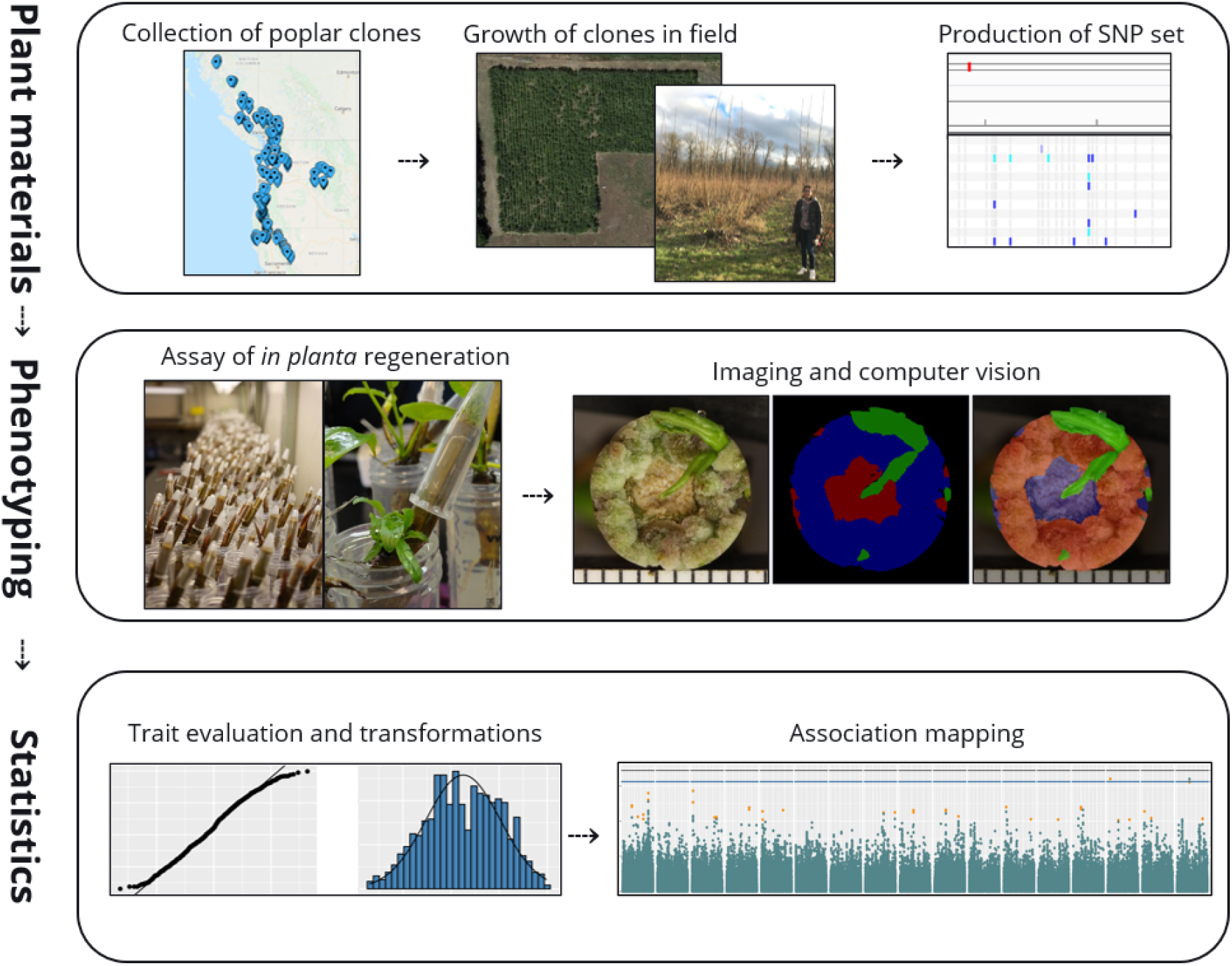
Overview of *in planta* callus and shoot regeneration GWAS workflow. The first row describes resources that were provided through other work prior to undertaking the GWAS. Images on the right of the “Phenotyping” panel are the RGB image, ground truth semantic label and overlay for one sample.

### Plant materials

We utilized an expanded version of the previously reported re-sequenced *P. trichocarpa* GWAS population (Zhang *et al*., 2018a; Tuskan *et al*., 2018; Muchero *et al*., 2018; Bdeir *et al*., 2019; Weighill *et al*., 2019; Chhetri *et al*., 2020). The population was recently expanded to include an additional 441 genotypes, filling a geographical gap that existed in first release with 882 genotypes (Fig. 2). While this clone bank is kept at multiple locations, phenotyping in this study only made use of the replicate in Corvallis, OR, featuring a total of 1,307 clones in the population (out of 1,323) and for 1,219 of which regeneration phenotyping was performed. Clones were grown at two field locations in Corvallis, OR: one location planted in 2009 featuring the original GWAS population, and another planted in 2015 featuring the newly added clones. Dormant cuttings were taken in the winters of 2018, 2019, and 2020, frozen, and then rooted up to one year later. Plants were then regularly pruned and fertilized to ensure there were healthy green leaves suitable for sterilization and introduction into tissue culture. A second greenhouse population was established and allowed to go dormant in winter; plants from this source were occasionally used to replace plants in the main greenhouse population that provided plant materials throughout the year.

**Figure 2.**
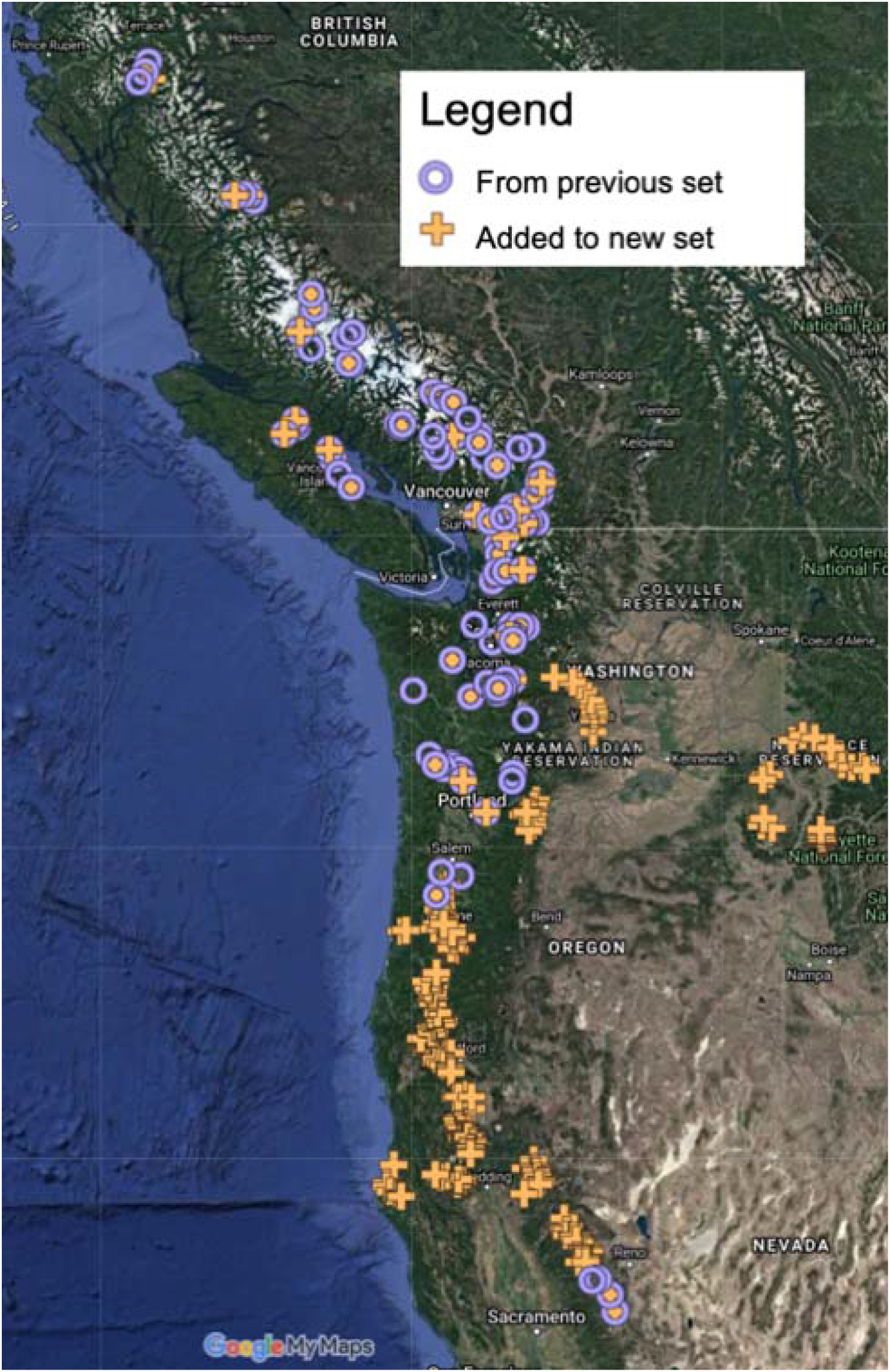
Origins of P. trichocarpa clones used to generate SNP set. A total of 1,323 native genotypes were collected over a geographical range across the pacific northwest region of the USA and the southwest of Canada. Tree location is shown for 1,301 genotypes for which precise location data is available.

### Sequencing and SNP set preparation

We performed SNP calling and analyzed the distribution of SNPs after resequencing of 406 additional genotypes by the DOE Joint Genome Institute, for a total of 1,323 genotypes in the complete SNP set (increased from 917 genotypes in the most recent SNP set). DNA short-read sequencing data for all 1,323 genotypes was filtered and trimmed with BBDuk (https://sourceforge.net/projects/bbmap/). Parameters used for BBDuk included ‘kmer=25’, ‘ktrim=r’, and ‘trimq=6’. Clean reads were then aligned to the P. trichocarpa v3.0 reference genome with bwa-mem v0.7.17 (Li & Durbin, 2009).

Duplicate reads were then marked with Sambamba markdup v0.7.0 (Tarasov *et al*., 2015). Next, BQSR was performed with GATK v4.1.3.0 BaseRecalibrator (Poplin *et al*., 2018). Variants were then called by sample with GATK v4.1.3.0 HaplotypeCaller and GVCFs were generated (Poplin *et al*., 2018). The GVCFs were then consolidated with GATK v4.1.3.0 GenomicsDBImport and joint calling was performed with GATK v4.1.3.0 GenotypeGVCFs. Next, SNPs and insertion-deletion variants (indels) were separated with GATK v4.1.3.0 SelectVariants. Indels were filtered with GATK v4.1.3.0 VariantFiltration with the following thresholds and flags: ‘-filter “QD < 2.0” --filter-name “QD2” -filter “FS > 200.0” --filter-name “FS200” -filter “ReadPosRankSum < −20.0” -- filter-name “ReadPosRankSum-20”’. SNPs were filtered with GATK v4.1.3.0 VariantFiltration with the following thresholds and flags, ‘-filter “QD < 2.0” --filter-name “QD2” -filter “SOR > 4.0” --filter-name “SOR4” -filter “FS > 60.0” --filter-name “FS60” -filter “MQ < 40.0” --filter-name “MQ40” -filter “MQRankSum < -12.5” --filter-name “MQRankSum-12.5” -filter “ReadPosRankSum < -8.0” --filter-name “ReadPosRankSum-8”’. There was a total of 40.4M SNPs prior to filtering for minor allele frequency (MAF) and additional quality criteria. The density and consistency of SNP data on each chromosome were assessed using the R package Cmplot (Fig. S1) and by producing histograms of gap sizes for each chromosome.

### Assay of regeneration

Frozen stem cuttings were incubated at 4 °C for 2-4 weeks, then placed in 50mL Falcon tubes with water for five weeks. Based on preliminary experiments (data not shown), we found that treatment of the decapitated surface of each stem with 100μL of 0.5mg/mL thidiazuron (TDZ) in water improved callus regeneration considerably (37% of genotypes produced shoots, compared to 24% without TDZ). After application of TDZ to a given stem tip, a 1.5mL microcentrifuge tube was inverted over the stem tip to prevent desiccation during regeneration (as shown in Fig. 1). On a weekly basis beginning the second week, decapitated stem surfaces were imaged from overhead using a Canon Rebel Xsi DSLR camera attached to a rack mount.

Due to practical limitations on the numbers of clones that could be assayed for regeneration simultaneously, subsets of the study genotypes (termed “phases”) were assayed at one time, with no more than 400 cuttings per phase. Images were taken on a weekly basis from the second week through the fifth week, with the exceptions of weeks four and five in the first phase and week four in the third phase. There were two replicate plants measured for all but the first three phases, where only a single replicate was used.

### Image segmentation

To perform annotation of images for computer vision, 249 images were randomly sampled from the first seven phases and manually annotated using the Intelligent Deep Annotator for Segmentation (IDEAS) graphical user interface that we previously developed (Yuan *et al*., 2022). As described in our prior work, these samples were used to train a convolutional neural network (PSPNet) via TensorFlow v1.14 to segment images of regenerating stem tips with each pixel labeled as one of four classes: callus, shoot, unregenerated stem and background. The trained model was deployed to produce inferred semantic masks for 11,791 images in the complete GWAS dataset, including 1,219 clones with up to four replicates and a median of two replicates, across four weekly timepoints (Fig. 3). At each timepoint, two traits were computed: the proportion of total plant area which consists of callus (henceforth, “callus area”), and of shoot (“shoot area”).

**Figure 3.**
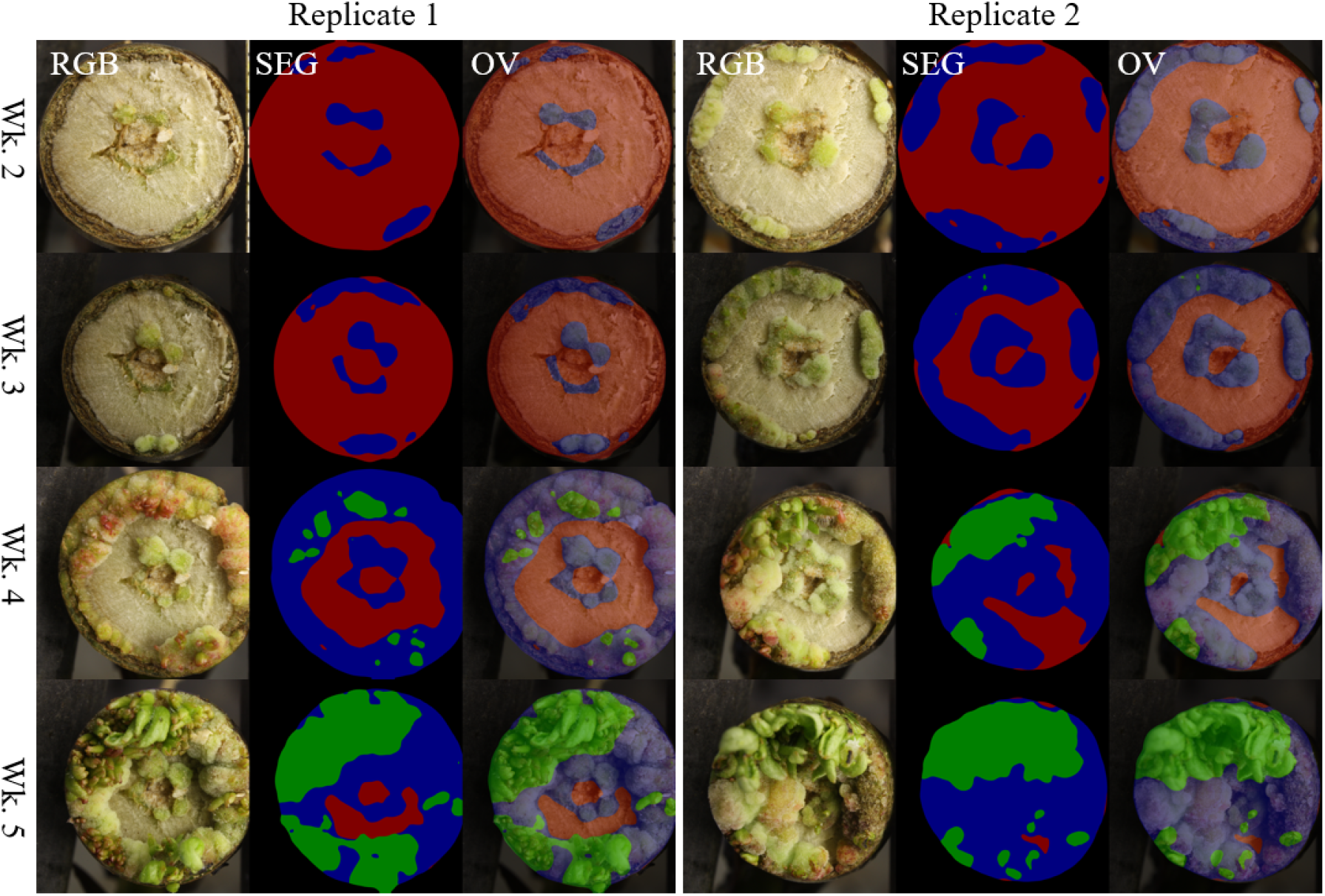
False-color examples of segmentation from regenerating stems shown in cross-section. For each section, shown are RGB images (left panels), inferred semantic mask labels (SEG, middle panels), and overlays of images and masks (OV, right panels) are shown for two replicates of genotype BESC-1013 across four timepoints of development, from week 2 (top) to week 5 (bottom). Semantic masks show unregenerated stem tissue (red), callus (blue) and shoot (green).

### Computer vision trait preparation for GWAS

For replicated samples, the mean value of each trait across the two replicates was computed and used in downstream analysis. For genotypes lacking replication due to infection in one replicate, the single unreplicated trait value was used.

Additional traits were computed by performing principal component analysis (PCA) using “stats::princomp” in R over three groups of traits: 1) callus area traits at all timepoints; 2) shoot area traits at all timepoints; and 3) both callus area and shoot area at all timepoints. Genotypes missing data for a given trait at any timepoint were excluded from a given PCA. Scree plots were evaluated to estimate the number of PCs representing significant proportions of trait variation.

The normality of traits was assessed using Q-Q plots, histograms, Shapiro-Wilks tests and Pearson correlation coefficients computed against theoretical normal distributions with the same mean and standard deviation as the given trait. To avoid severe violations of normality that may lead to inflated error rates, all traits were transformed prior to GWAS methods that depend on normality and linear model assumptions. The most basic transformation applied was a removal of zero values followed by Box-Cox transformation. For certain PC traits, a spike was observed at particular values, which corresponded to genotypes with zero values for all traits used in the given PCA; these genotypes were consequently removed. In cases where we determined that thresholding or extreme outlier removal was necessary, these treatments were performed prior to the Box-Cox transformation. In addition, as an alternative to Box-Cox transformations, rank-based inverse normal (RB-INV) transformations were performed for difficult distributions (Fig. S2, Table S1, S2).

### Association Mapping

Because of the distinct assumptions and data types for which various GWAS methods are suited, along with the non-normality of computer-vision-based traits and the risk of sacrificing statistical power with transformations, we employed an analysis pipeline that made use of four GWAS methods. These include Genome-wide Efficient Mixed Model Association (GEMMA; Zhou & Stephens, 2012), which was parallelized using the GNU Parallel (Tange, 2020) framework to simultaneously run each given trait on a CPU core. Due to an assumption of normally distributed model predictions for GEMMA, this method was only used to study versions of traits that had been transformed toward normality, including the exclusion of all samples with zero values. The Generalized Mixed Model Association Test (GMMAT; Chen *et al*., 2016b) was used for single-marker tests with the same kinship covariate as GEMMA; however, rather than using continuous trait variables, GMMAT applies logistic regression and works with binarized traits. Finally, as a means of performing tests that avoid any transformation or binarization of traits, we applied our Multi-Threaded Monte Carlo Sequence Kernel Association Test (MTMC-SKAT; <u>github.com/naglemi/mtmcskat</u>), an R extension of the established SKAT method (Wu *et al*., 2011; Ionita-Laza *et al*., 2013) extending support for efficient resampling on high-performance clusters. Using MTMC-SKAT, we tested for the combined effect of adjacent SNPs within 3kb windows and computed empirical *p*-values for SNPs indicated by an initial parametric test as being strongly associated with a trait. With MTMC-SKAT, we tested and compared the control for population structure via two types of covariates: a matrix of PCs from PCA of SNP data (P model) and a matrix of theoretical subpopulation ancestry estimates from fastSTRUCTURE (Q model; Raj *et al*., 2014). MTMC-SKAT was deployed on the COMET high-performance cluster made available through the NSF XSEDE program (Towns *et al*., 2014). Additional details on GWAS methods can be found in Supplementary Methods.

Statistically significant associations from the various pipelines were first determined by computing Benjamini-Hochberg false discovery rate (FDR; α = 0.10) and Bonferroni thresholds (α = 0.05). SNPs not found on contiguous assembled chromosomes were excluded from computation from this point forward (approximately 3.6% of total SNPs). The Bonferroni thresholds were computed given the number of tests equal to the number of filtered SNPs (for single-marker tests GEMMA and GMMAT) and the number of 3kb SNP windows (in the case of SKAT). We then extracted lists of SNPs with *p*-values below these thresholds for interrogation.

We next evaluated the extent to which multiple SNPs supported the association of a nearby gene, whether individual SNPs met the FDR or Bonferroni statistical thresholds or not. We implemented the Augmented Rank Truncation (ART) method (Vsevolozhskaya *et al*., 2019) to scan Wald *p*-values from GEMMA and GMMAT and identify cases where a SNP produces a *p*-value below 1*10^-5^ and is within 500bp of at least 5 additional SNPs with *p*-values below 1*10^-4^ when considering the upper half of top-ranking SNPs (according to *p*-values). While we initially computed Wald *p-*values with GMMAT for only the top 100 or 1,000 SNPs according to a score test, additional Wald *p-*values were computed for use in ART. For each of these 1kb windows, a combined *p*-value was computed for the extracted SNPs. A Bonferroni threshold for ART *p*-values was computed (α = 0.05) from the approximate number of independent tests (contiguous assembled genome size / ART window size). The Bonferroni threshold of ∼1.27*10^-7^ was computed using the number of independent tests of (∼3.94*10^6^ 1kb windows spanning the ∼394 Mb of contiguous assembled chromosomes) and is notably less conservative than the Bonferroni threshold used for raw *p*-values from GEMMA/GMMAT (henceforth, “conservative Bonferroni”), as computed from the total number of tests (as low as ∼4*10^-9^, given up to ∼13 million SNPs on contiguous chromosomes remaining after filtering; Supplementary Methods). To be emphasized, it is well-known that Bonferroni thresholds for individual SNPs consider each SNP as an independent test, leading to overcorrection since large numbers of SNPs are in LD.

To determine on a high-throughput scale which genes are likely to be responsible for statistically significant quantitative trait loci (QTLs; either SNPs or SNP windows), we used R scripts to reference genome and genome annotation data available through Phytozome (phytozome.jgi.doe.gov; Tuskan *et al*., 2006). In this workflow, the positions of loci were evaluated for candidate genes only when these loci represent the “peak” of a signal, determined by checking for any other loci within 30kb with a more significant *p*-value. The candidate gene responsible for the significance of a given locus was assumed by the workflow to be the gene that encompasses or is closest to the locus. The R package InterMineR (Kyritsis *et al*., 2019) was used to collect Phytozome data on gene function, Arabidopsis homologs, and gene ontology terms and organized these by locus. The GreeNC database was used to identify possible noncoding regulatory RNAs among gene candidates (Di Marsico *et al*., 2022). For the top gene candidates, particularly those passing the conservative Bonferroni or FDR (α = 0.10) thresholds, or those passing the less conservative Bonferroni thresholds used for ART and among the five most-significant GEMMA-ART or GMMAT-ART associations for a given trait, Integrative Genomics Viewer (IGV; Robinson *et al*., 2011; Thorvaldsdóttir *et al*., 2013) was used to manually investigate gene position relative to significant SNPs, including consideration of other nearby genes, distance to the putative transcription start site, and direction of transcription.

## Results

### Trait transformations

Prior to transformations, most traits displayed marked non-normal characteristics as indicated by Q-Q plots, histograms, Shapiro-wilk tests, and Pearson correlation coefficients of each distribution with a normal distribution with the same mean and standard deviation. The improvement in normality after transformation was marked in most cases (Table S1, S2; Fig. S2; data not shown). Non-normal characteristics of most traits were reduced substantially by excluding genotypes with zero values and applying a Box-Cox transformation (e.g., Fig. S2). For the traits of callus or shoot area at each timepoint, based on visual inspection of histograms and Q-Q plots, the improvement in metrics of normality was deemed adequate for linear models. All PCA-derived traits necessitated additional treatments to avoid severe violations of the normality assumption of linear models, including removal of outliers and in some cases removal of values below an elbow in the frequency distribution (estimated as the position where the second derivative of the probability frequency distribution is maximum; Table S2).

### Principal components as proxies for complex patterns of regeneration

Scree plots and heat maps of loadings revealed common trends in regeneration across timepoints and regenerating tissue types (callus and shoot). These results were obtained for three different sets of PCA with different groups of traits: first, for both callus and shoot area at all timepoints (Fig. 4), and then with callus and shoot data analyzed independently over all timepoints (Fig. S3). In all three cases, the PC explaining the most variation (PC1) represented a tendency of the tissue(s) included in PCA to regenerate across all timepoints. Latter PCs provided proxies for more complex patterns of regeneration. PC2 over callus traits appears to represent high levels of callus regeneration at early, but not later timepoints. PC2 over all callus and shoot traits appears to represent a tendency for callus to regenerate robustly, but to fail to develop into shoots. Subsequent PCs, for each batch of traits, represented a relatively small proportion of variance explained and were thus not analyzed for gene candidates.

**Figure 4.**
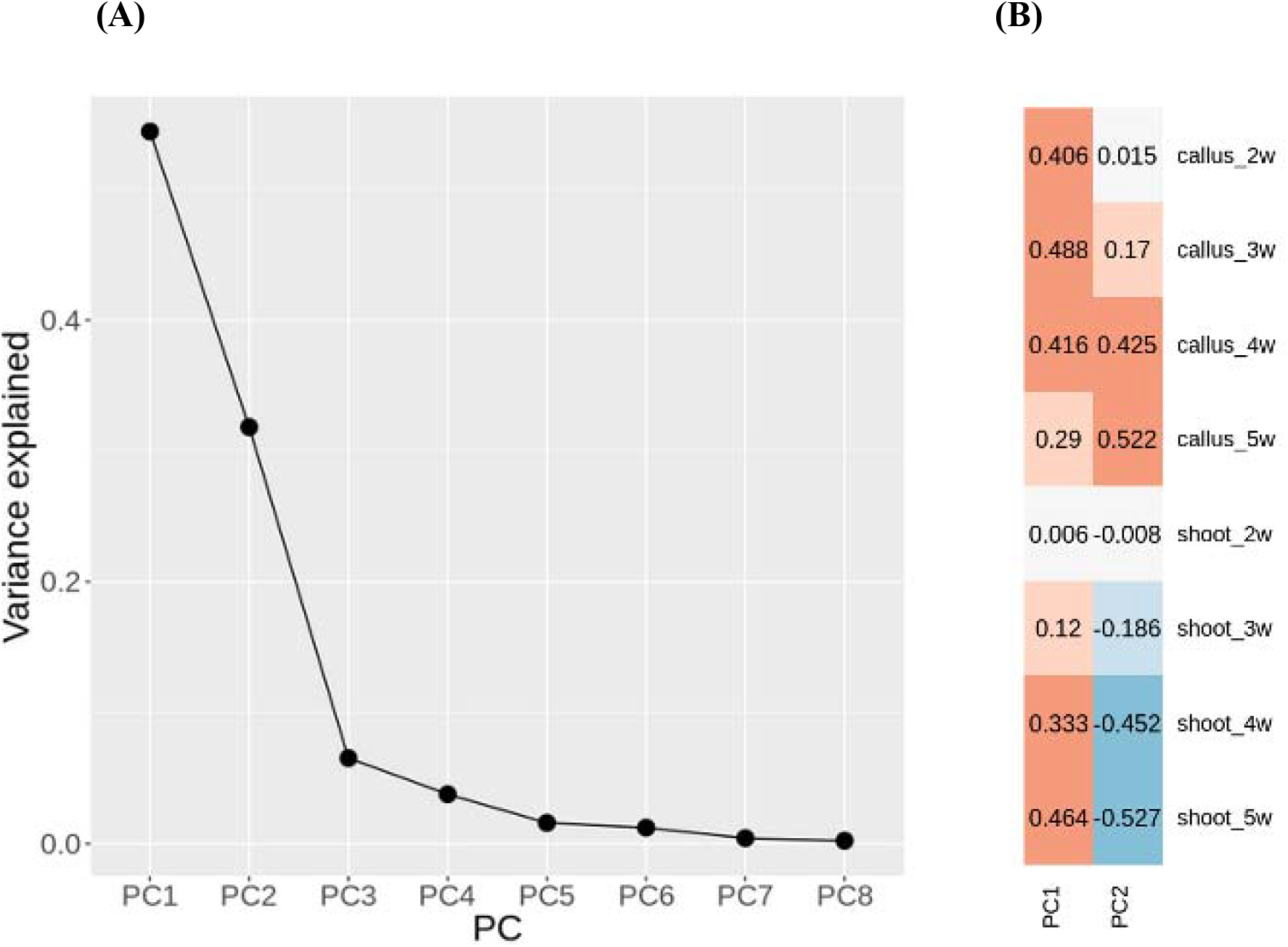
PCA of *in planta* callus and shoot regeneration traits. Results are shown from PCA over all callus and shoot traits across four weekly timepoints. **(A)** Scree plot; **(B)** Heat map of loadings from PCA.

### SNP set for *P. trichocarpa* provides comprehensive view of natural variation

The SNP set produced for this population displays polymorphism across all regions of contiguous chromosomes represented in the reference genome (Fig. S1). Poplar clones collected for the GWAS clone bank represent a wide range of geographic diversity, nearly spanning the natural range of *P. trichocarpa* from British Columbia to the Pacific Northwest of the United States (Fig. 2). There is clearly very strong natural intrachromosomal recombination, as LD decay occurs rapidly and reaches R^2^ = 0.2 within 2kb whether computed for the whole population or either of the two most prominent subpopulations (Fig. 5, Fig. S4). Population structure analysis with fastSTRUCTURE, cross-referenced with geography and phylogenetics, supported the existence of 6 or 7 ancestral subpopulations, which were most distinct in southern regions, while a greater degree of admixture was prevalent farther north and may be responsible for the significantly lower LD in the “Oregon” subpopulation (P < 0.001; Fig. S4). Our linear models supported a positive relationship between latitude and callus traits, but were unable to detect such a relationship for shoot traits (Fig. S5-S10; Table S3; Supplemental Results and Discussion).

**Figure 5.**
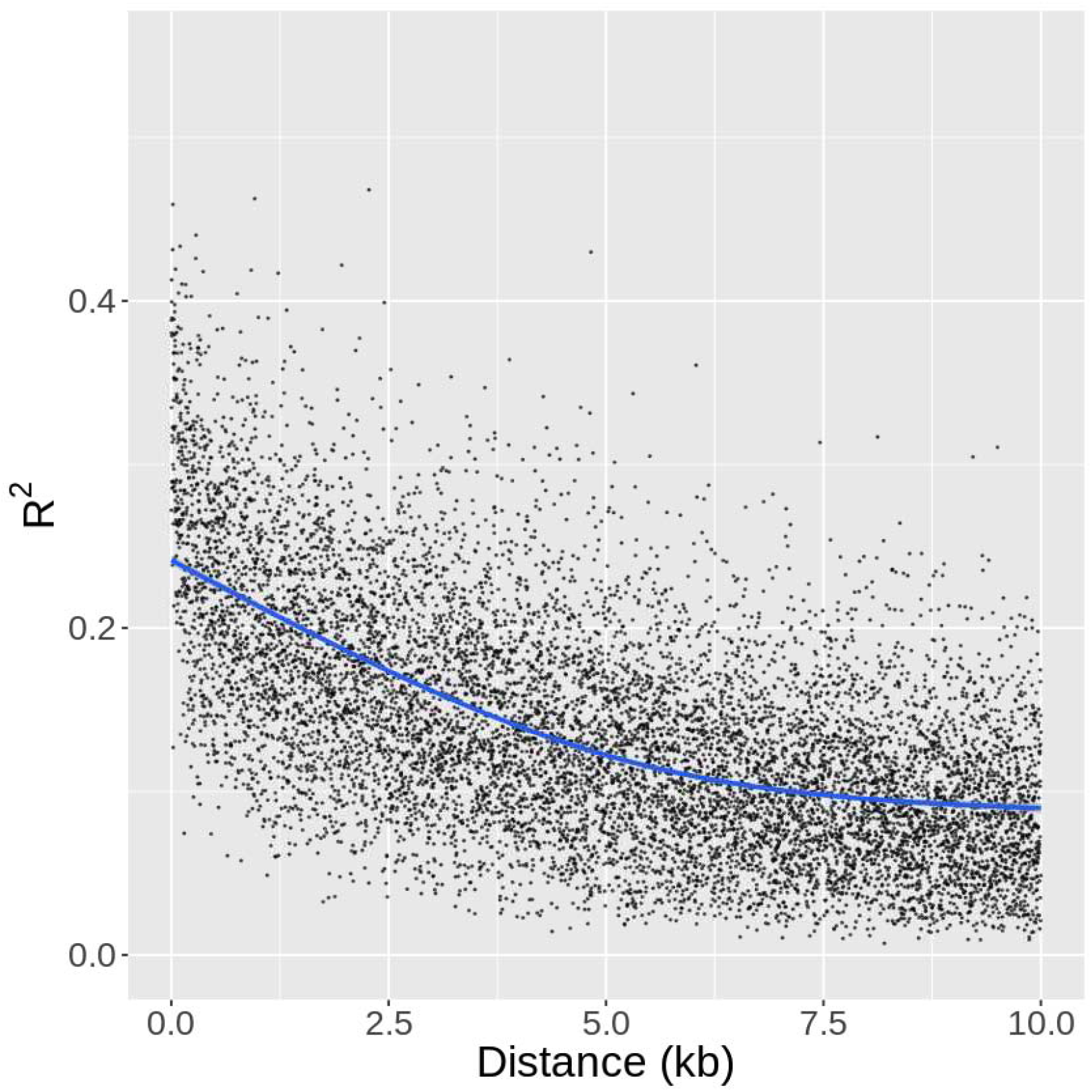
Linkage disequilibrium decay. Each point represents the mean LD between SNPs of a given distance (x-axis). The blue line represents LD decay as computed with a spline function.

### Genes implicated by significant quantitative trait loci (QTLs)

We interrogated traits with *h^2^_SNP_* above 0.10 for candidate genes (Table S4), except for shoot area at week 2 (first timepoint) due to a sparse distribution and high level of noise for this trait. We utilized three multiple-testing correction methods (from most to least conservative: conservative Bonferroni, Benjamini-Hochberg FDR and ART-Bonferroni) to provide three levels of confidence in associations to support prioritization of gene candidates. Across the three GWAS methods GEMMA, GMMAT, and (MTMC-)SKAT, we report a total of 13 unique QTL peaks with *p*-values passing the conservative Bonferroni significance threshold, as well as 44 passing the FDR (α = 0.10) threshold. Among Bonferroni-significant associations, 6 are inside or within 5kb of a gene found in the genome annotation, as well as 32 associations (∼72.7%) meeting the FDR threshold (Fig. 6-8, Table S5, S6). We found 137 unique QTL peaks from applying our implementation of ART to GEMMA results (Table S5, S7), as well as 24 from applying ART to GMMAT results (Table S5, S8). Several of the most promising candidates, based on the biology of their homologs in Arabidopsis, are shown in Table 1.

**Figure 6.**
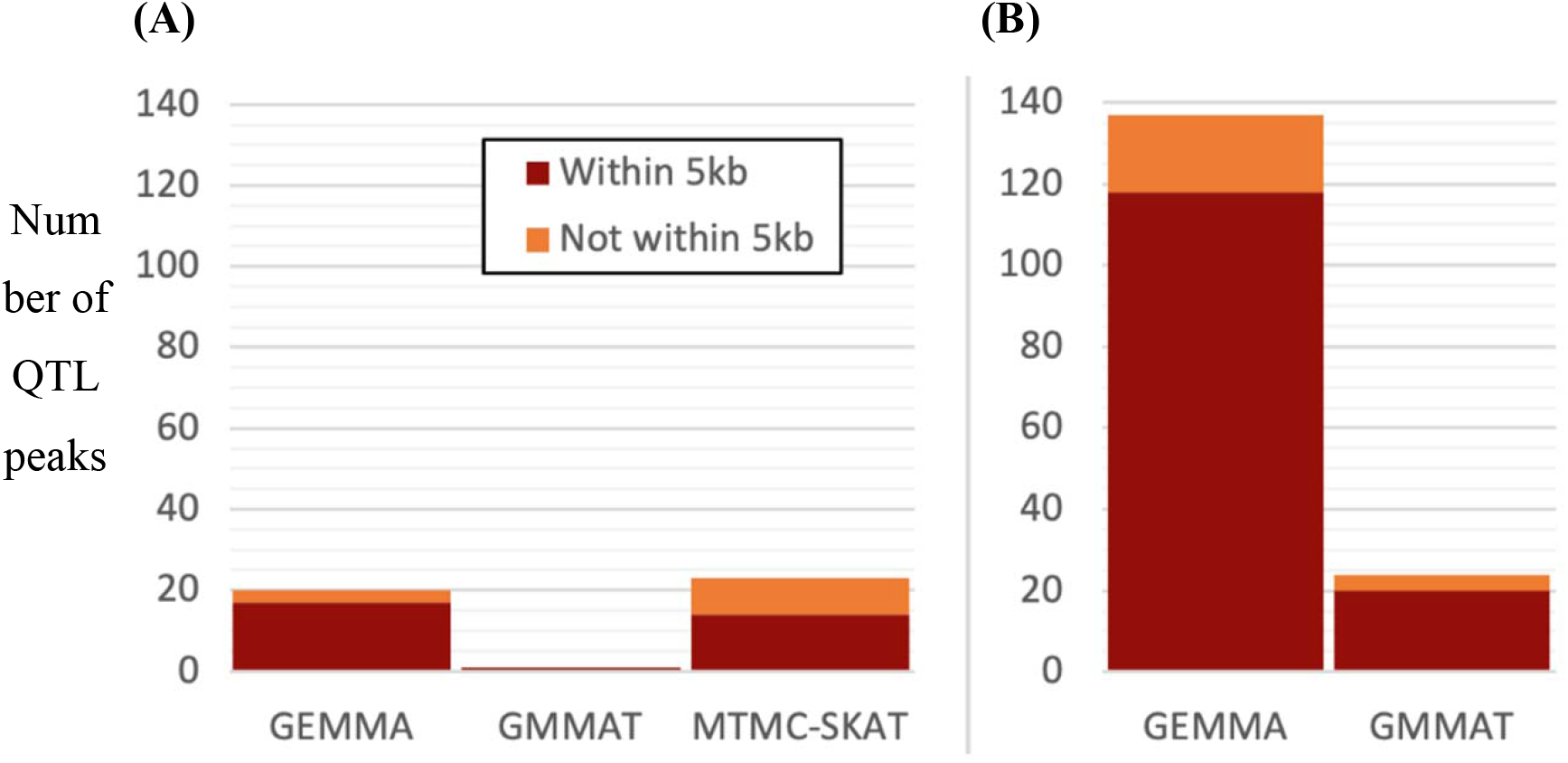
Tallies of *in planta* callus and shoot associations across GWAS methods. Shown are barplots summarizing the numbers of associations from each GWAS method, with two types of significance thresholds, as well as within a 5kb distance threshold of the nearest gene. QTL peaks were taken as the point with the lowest *p*-value at any given peak, where multiple points within the same peak may otherwise pass a given significance threshold. **(A)** QTL peaks passing the Benjamini-Hochberg threshold (FDR; α = 0.10) and/or conservative Bonferroni threshold; **(B)** QTL peaks passing ART-Bonferroni threshold (α = 0.05, *N* of 1kb windows in genome).

**Figure 7.**
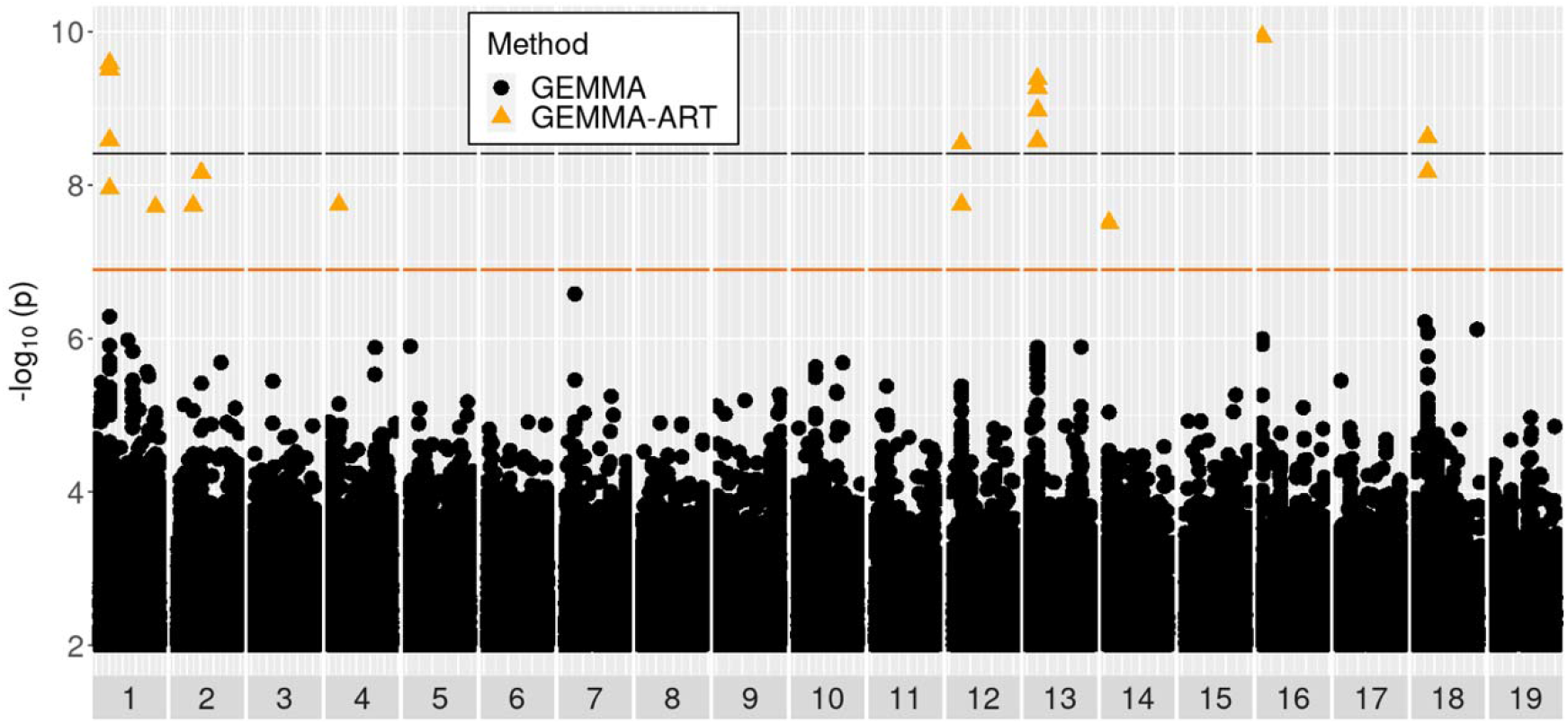
Manhattan plot of GEMMA-ART results for week four callus area. Shown is a Manhattan plot for GEMMA results for the trait of callus area at week four. Black and orange lines show Bonferroni significance thresholds for GEMMA results with independent SNPs, and for ART applied to GEMMA over 1kb windows of SNPs, respectively. Black circles represent tests of individual SNPs by GEMMA, while orange triangles represent 1kb windows tested by ART applied to GEMMA results.

**Figure 8.**
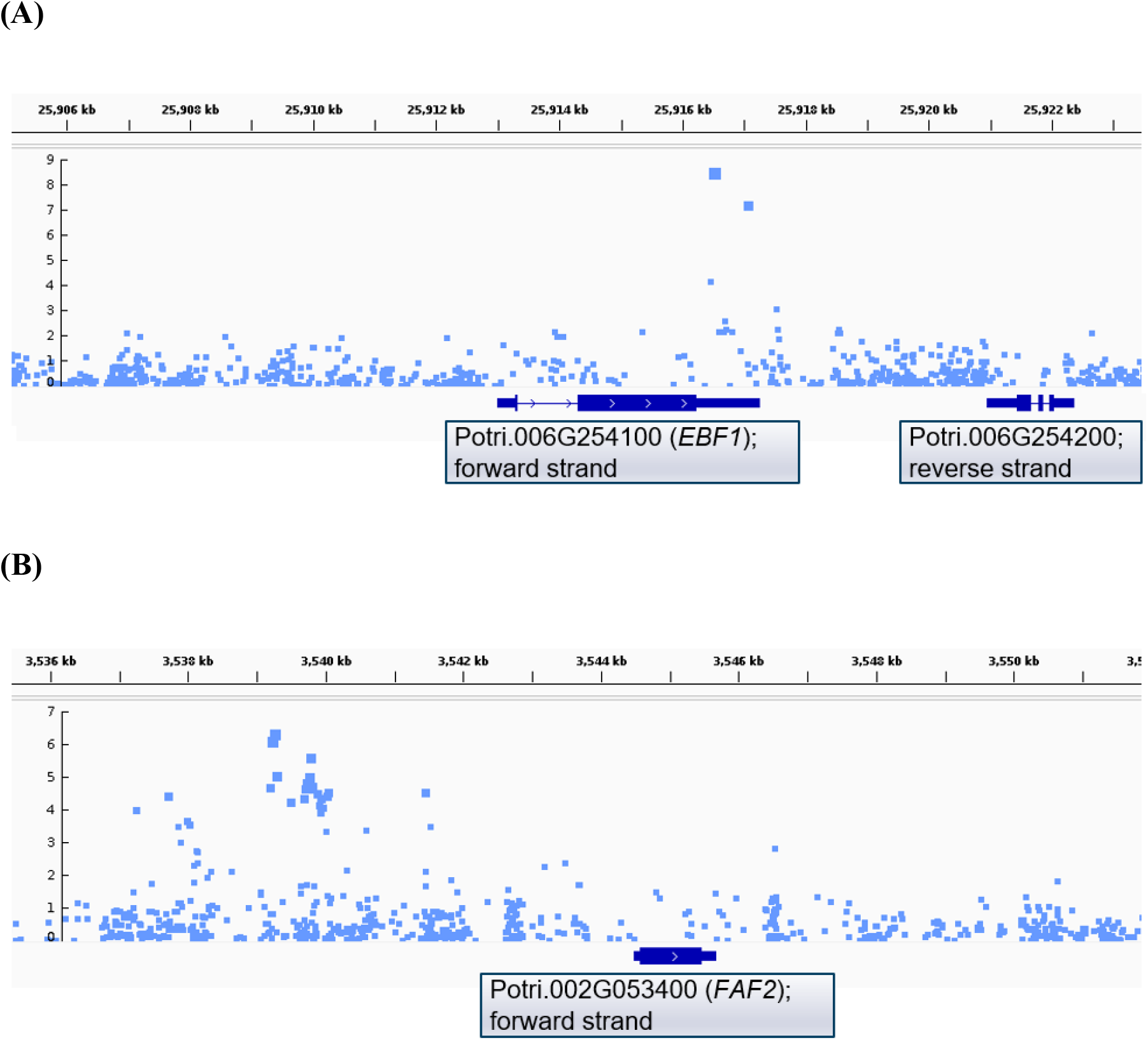
Close-up view of Manhattan plots for example *in planta* regeneration traits aligned with genome annotation. Shown are show zoomed-in portions of Manhattan plots aligned to the genome annotation for P. trichocarpa (v3.1). Introns, untranslated regions and exons are respectively visualized with increasing thickness of bars. Labels in gray boxes were manually added to show gene IDs and the strand on which genes are found. **(A)** Results on chromosome 6 for GEMMA of Box-Cox-transformed trait Shoot PC2; **(B)** Results on chromosome 2 for GEMMA of Box-Cox transformed trait Callus/Shoot PC1, showing an association found significant via ART. These plots were made with Integrative Genomics Viewer (IGV) and boxes with Accession ID, gene name and strand were manually added underneath. Examples of plots for additional loci can be found in Fig. S14.

**Table 1.**
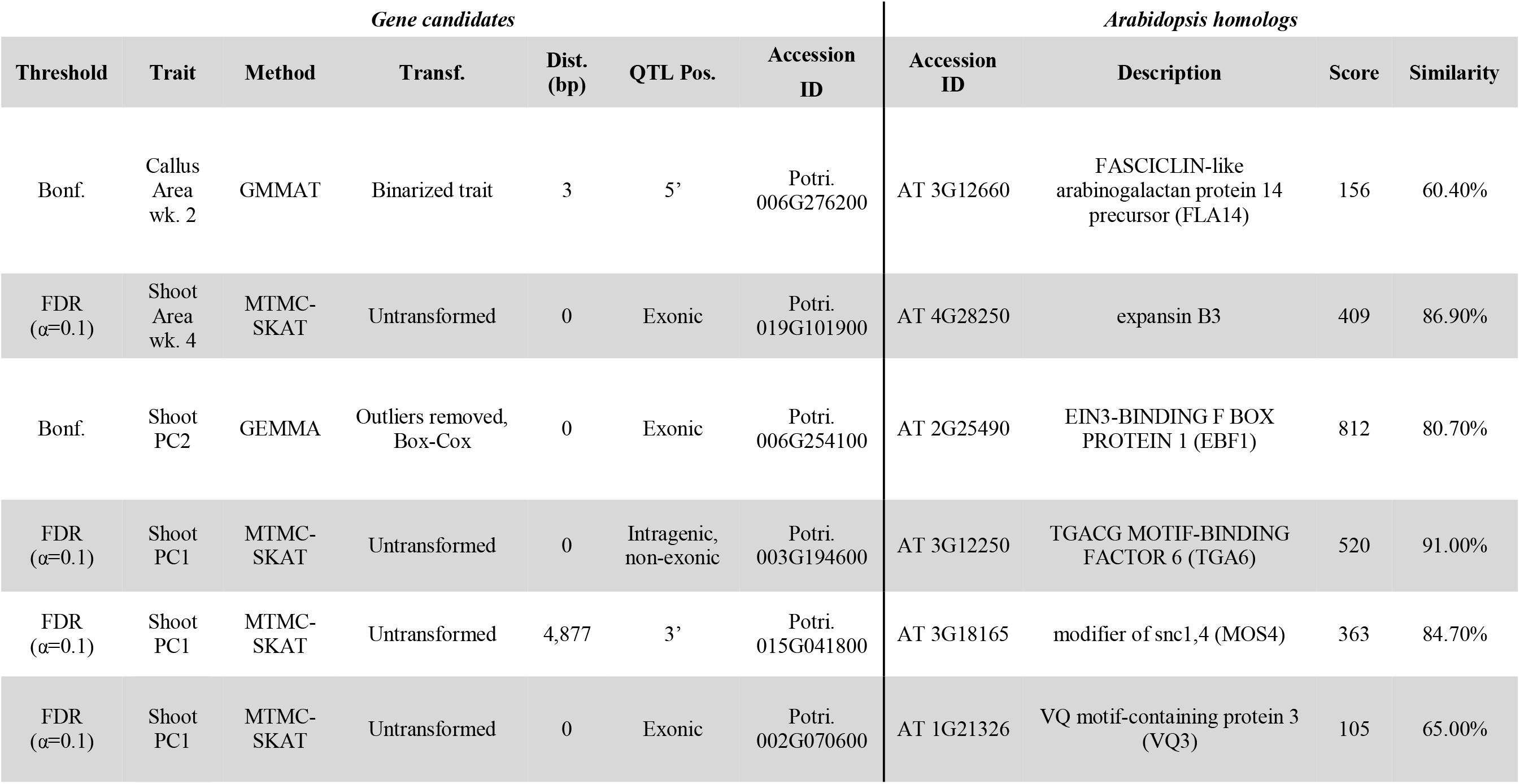
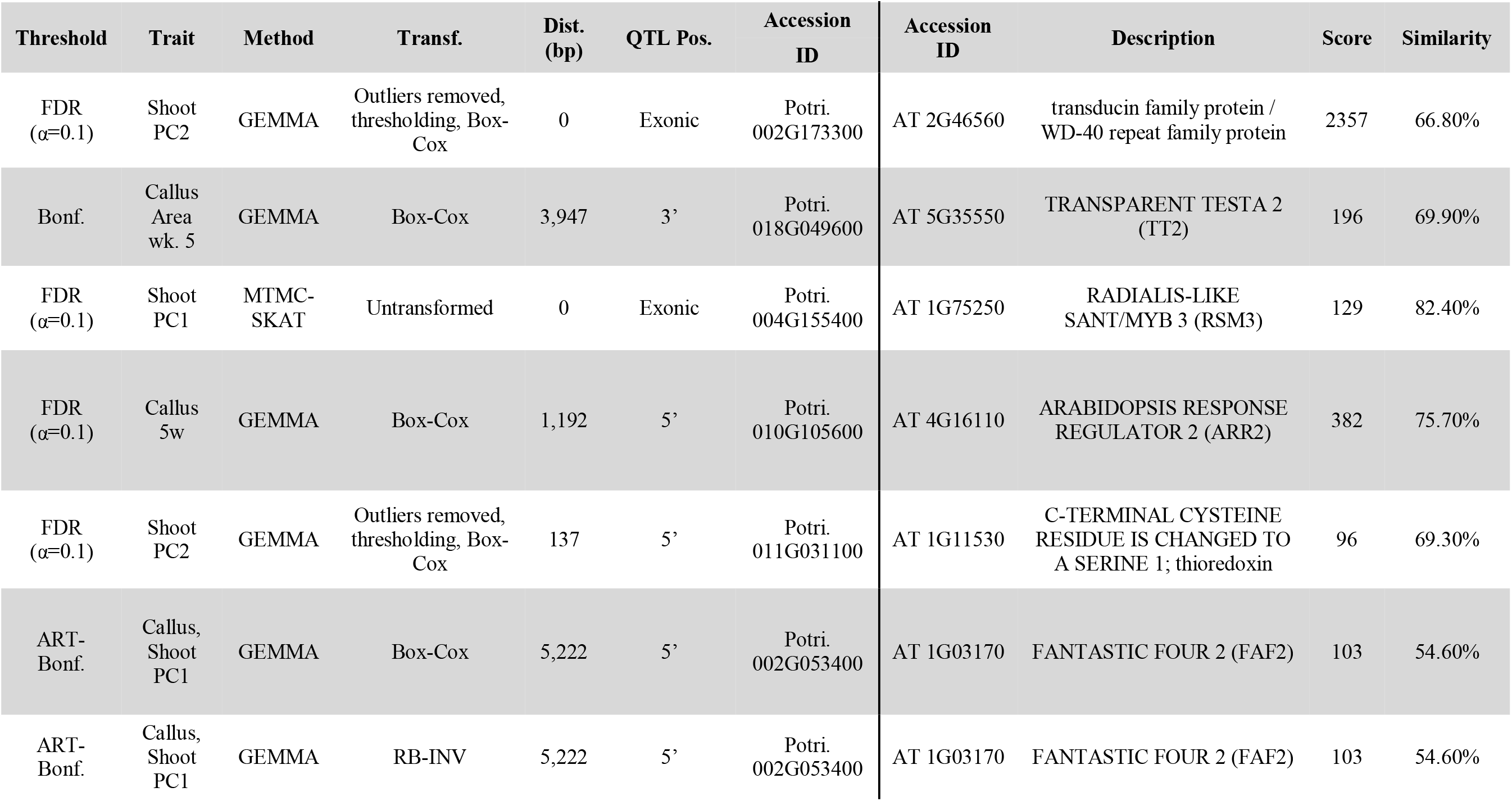
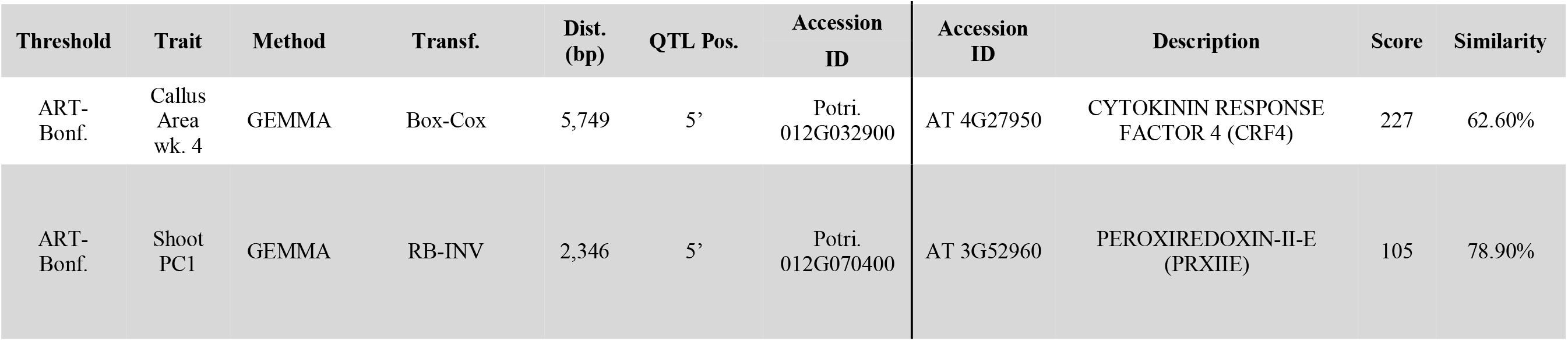
Fourteen *in planta* regeneration GWAS candidates highlighted for discussion, and Arabidopsis homologs relevant to regeneration. Relevant literature is discussed for each of these candidates (Discussion). Distance (Dist.) of individual SNPs or SKAT/ART window centers from the transcription start site is shown for intergenic associations. Score and similarity percentage is shown for Smith-Waterman alignment of poplar gene candidates with Arabidopsis homologs, extracted from Phytozome (phytozome.jgi.doe.gov). Remaining gene candidates are summarized in Table S6-8.

| *Gene candidates* |  |  |  |  |  |  | *Arabidopsis homologs* |  |  |  |
| --- | --- | --- | --- | --- | --- | --- | --- | --- | --- | --- |
| Threshold | Trait | Method | Transf. | Dist. (bp) | QTL Pos. | Accession ID | Accession ID | Description | Score | Similarity |
| Bonf. | Callus Area wk. 2 | GMMAT | Binarized trait | 3 | 5' | Potri. 006G276200 | AT 3G12660 | FASCICLIN-like arabinogalactan protein 14 precursor (FLA14) | 156 | 60.40% |
| FDR ($\alpha=0.1$) | Shoot Area wk. 4 | MTMC-SKAT | Untransformed | 0 | Exonic | Potri. 019G101900 | AT 4G28250 | expansin B3 | 409 | 86.90% |
| Bonf. | Shoot PC2 | GEMMA | Outliers removed, Box-Cox | 0 | Exonic | Potri. 006G254100 | AT 2G25490 | EIN3-BINDING F BOX PROTEIN 1 (EBF1) | 812 | 80.70% |
| FDR ($\alpha=0.1$) | Shoot PC1 | MTMC-SKAT | Untransformed | 0 | Intragenic, non-exonic | Potri. 003G194600 | AT 3G12250 | TGACG MOTIF-BINDING FACTOR 6 (TGA6) | 520 | 91.00% |
| FDR ($\alpha=0.1$) | Shoot PC1 | MTMC-SKAT | Untransformed | 4,877 | 3' | Potri. 015G041800 | AT 3G18165 | modifier of snc1,4 (MOS4) | 363 | 84.70% |
| FDR ($\alpha=0.1$) | Shoot PC1 | MTMC-SKAT | Untransformed | 0 | Exonic | Potri. 002G070600 | AT 1G21326 | VQ motif-containing protein 3 (VQ3) | 105 | 65.00% |
| Threshold | Trait | Method | Transf. | Dist. (bp) | QTL Pos. | Accession ID | Accession ID | Description | Score | Similarity |
| FDR ($\alpha=0.1$) | Shoot PC2 | GEMMA | Outliers removed, thresholding, Box-Cox | 0 | Exonic | Potri. 002G173300 | AT 2G46560 | transducin family protein / WD-40 repeat family protein | 2357 | 66.80% |
| Bonf. | Callus Area wk. 5 | GEMMA | Box-Cox | 3,947 | 3' | Potri. 018G049600 | AT 5G35550 | TRANSPARENT TESTA 2 (TT2) | 196 | 69.90% |
| FDR ($\alpha=0.1$) | Shoot PC1 | MTMC-SKAT | Untransformed | 0 | Exonic | Potri. 004G155400 | AT 1G75250 | RADIALIS-LIKE SANT/MYB 3 (RSM3) | 129 | 82.40% |
| FDR ($\alpha=0.1$) | Callus 5w | GEMMA | Box-Cox | 1,192 | 5' | Potri. 010G105600 | AT 4G16110 | ARABIDOPSIS RESPONSE REGULATOR 2 (ARR2) | 382 | 75.70% |
| FDR ($\alpha=0.1$) | Shoot PC2 | GEMMA | Outliers removed, thresholding, Box-Cox | 137 | 5' | Potri. 011G031100 | AT 1G11530 | C-TERMINAL CYSTEINE RESIDUE IS CHANGED TO A SERINE 1; thioredoxin | 96 | 69.30% |
| ART-Bonf. | Callus, Shoot PC1 | GEMMA | Box-Cox | 5,222 | 5' | Potri. 002G053400 | AT 1G03170 | FANTASTIC FOUR 2 (FAF2) | 103 | 54.60% |
| ART-Bonf. | Callus, Shoot PC1 | GEMMA | RB-INV | 5,222 | 5' | Potri. 002G053400 | AT 1G03170 | FANTASTIC FOUR 2 (FAF2) | 103 | 54.60% |
| Threshold | Trait | Method | Transf. | Dist.<br>(bp) | QTL Pos. | Accession<br>ID | Accession<br>ID | Description | Score | Similarity |
| ART-Bonf. | Callus<br>Area<br>wk. 4 | GEMMA | Box-Cox | 5,749 | 5' | Potri.<br>012G032900 | AT 4G27950 | CYTOKININ RESPONSE<br>FACTOR 4 (CRF4) | 227 | 62.60% |
| ART-Bonf. | Shoot<br>PC1 | GEMMA | RB-INV | 2,346 | 5' | Potri.<br>012G070400 | AT 3G52960 | PEROXIREDOXIN-II-E<br>(PRXIIE) | 105 | 78.90% |

## Discussion

### High-throughput phenomics support scale and precision of GWAS

Our mapping of complex developmental traits relied on computer vision methods to precisely quantify traits that would be impractical to measure manually. Although the use of an ordinal scoring system instead of segmentation is an alternative approach, this would have risked the introduction of subjective biases and violation of linear model assumptions—while sacrificing much precision and detail. The high-throughput phenomics workflow used for this work was described, in part, by Yuan et al. (2022). The IDEAS graphical interface for image annotation enabled the production of a large set of training and validation examples (249 images in total) with pixelwise labels for callus, shoot and unregenerated tissues. Here, we applied this method on a high-throughput scale for GWAS and evaluated statistical consequences and necessary downstream adaptations, such as appropriate trait transformations and GWAS methods. This training set enabled a deep segmentation model that was used to automatically segment a total of 11,791 images. Although generation of the training samples was time-consuming, performing manual segmentation for all images would have been time-prohibitive. Image annotation using the GUI required approximately five minutes per image using IDEAS, for a total of approximately 21 hours. Without a trained model for image segmentation, annotation of the whole dataset would have required nearly 1,000 hours. This system or others that are functionally comparable (Russell *et al*., 2008; Dutta & Zisserman, 2019) can be made more accessible and practical with innovations to reduce the number of clicks needed for image annotation by further semi-automation of annotation. While there have been many successes in the use of color thresholding methods that segment plants while requiring relatively little or no training data (Yang *et al*., 2015; Das *et al*., 2015; Al-Tamimi *et al*., 2016; Guo *et al*., 2018; Gage *et al*., 2018; Pham *et al*., 2019; Seethepalli *et al*., 2020; Campbell *et al*., 2020; Ogawa *et al*., 2021; Affortit *et al*., 2022), the highly variable colors observed in callus and shoot rendered these methods inappropriate for our study. Furthermore, in our previous work we reported a comparison of Random Forests (RF) to our trained PSPNet model and found RF to underperform in segmentation accuracy (Yuan *et al*., 2022).

As reported in our prior work (Yuan *et al*., 2022), overall accuracy in segmentation of the “validation” set of images was 79.21% as measured by IoU, while relatively homogenous stem tissues had IoU of 88.14%, and highly heterogenous callus tissues had 67.40%. Imperfections in defining segmentation boundaries, whether manually or using a trained model, leads to the introduction of statistical noise that will tend to reduce the computed heritability of traits. However, the use of PCA over batches of traits reduced the impact of this noise to produce traits of greater heritability. Indeed, while we studied six PC traits and eight non-PC traits, four of the six traits with the highest heritability with were PCs (Table S4).

Clearly, there is room for further advancement to make these phenomic workflows more efficient and user-friendly. Advances in the architectures of deep segmentation neural networks can contribute to improved accuracy in segmenting complex and heterogenous tissues of interest to biologists. Further innovation in the interfaces used to generate training sets can reduce the time cost needed for training these models. Once a training set is obtained, the training of neural networks can require a labor-intensive process of hyperparameter tuning. Recent developments such as Automated Machine Learning (AutoML) can be used to automate the hyperparameter tuning process and simplify training (Waring *et al*., 2020; Karmaker *et al*., 2021). Finally, the continued development of robust and efficient GWAS methods that avoid linear model assumptions can help to maximize the genetic insights gained from non-normal traits based on image segmentation.

### Complementary GWAS approaches provide variety of insights

Transformation of traits to approximate normality is commonly employed for biological data during GWAS to avoid violating linear model assumption of normality of residuals. In our study, because traits were computed as the proportion of plant tissue labeled by segmentation masks as callus or shoot, and many genotypes did not produce either tissue at early timepoints or any timepoints, the resulting distributions featured a mix of a zero and nonzero values. Among traits in our study, the proportion of genotypes with zero values ranged from 89 (for callus area at week five) to 1,106 (for shoot area at week two). To avoid violations of the normality assumption arising from noncontinuous distributions, genotypes featuring zero values were excluded from GEMMA tests for each trait, but presence/absence tests were performed using GMMAT. GMMAT and SKAT offer two complementary approaches to avoid problematic assumptions without trait transformation, thus obviating the need to exclude totally recalcitrant genotypes and thus suffer reduced statistical power.

Single-SNP methods including GEMMA and GMMAT share the advantage of providing insights into the specific SNPs most likely to be causative with respect to the effect of a gene on a trait. In most cases in our results, these appear to be regulatory SNPs in promoters, suggesting that variation in gene expression, rather than sequence, is the primary cause of trait variation. However, single-SNP methods suffer from relatively low statistical power since by their nature they treat each SNP-trait relationship as an independent test and do not consider combined effects of nearby SNPs. In contrast, SKAT provides improved statistical power by allowing tests for the combined effects of adjacent SNPs grouped into SNP windows, but only provides a single *p*-value for a whole SNP window. Thus, our SKAT results do not make clear which SNPs in a given window are responsible for trait variation, and as windows often overlap coding and regulatory regions, we lack insight into whether SKAT-implicated candidates are responsible for trait differences due to variation in their regulation or protein structure. Moreover, even when a given window is entirely intergenic, we lack an ability for straightforward investigation of specific promoter motifs that may be implicated by SKAT due to the lack of single-SNP resolution. Finally, SKAT involves the upweighting of rare SNPs and results are therefore less likely to feature top gene candidates regulated by common variation (Wu *et al*., 2011; Ionita-Laza *et al*., 2013).

We therefore sought to employ a “best of both worlds” approach to improve the statistical power of GEMMA and GMMAT by considering combined effects of adjacent common SNPs without losing clarity into the specific SNPs most likely to be causative. To this end, we employed ART as a post-hoc analysis of GEMMA and GMMAT results. As ART involves the computation of combined *p*-values over SNP windows and does not assume independence of SNPs, we obtained an increase in statistical power both via both reduced *p*-values for SNP windows compared to individual SNPs (Vsevolozhskaya *et al*., 2019), and by the ability to use a less-stringent Bonferroni threshold due to the number of tests being equal to the number of 1kb SNP windows rather than the number of individual SNPs. Our usage of ART enabled the detection of candidate genes including *FAF2*, *CRF4* and *PRXIIE* (Table 1) that otherwise would have been missed in our study. Although we are unaware of applied GWAS studies utilizing ART, our results demonstrate the potential for this method to increase effective statistical power in GWAS, which may be of particular benefit to studies using segmentation-based traits that follow non-normal distributions and undergo transformations with a risk of reducing power. As previously mentioned (Results), the ART-Bonferroni threshold represents the least conservative of our three multiple-testing correction thresholds; associations meeting this threshold alone thus represent our lowest-priority associations for investigation and discussion.

### Candidate genes have diverse roles in signaling and development

Our results indicate that natural variation in capabilities for *in planta* regeneration in poplar is controlled by numerous genes with functionally diverse roles, including in cell wall and membrane structure, hormone signaling, anthocyanin production and reactive oxygen species (ROS) regulation. Very few gene of the gene candidates we found were previously reported in two previous GWAS of related poplar traits, including several in a GWAS of adventitious shoot and root in hydroponic culture (Sun *et al*., 2019) and none in a GWAS of *in vitro* callus regeneration (Tuskan *et al*., 2018), likely due in part to the different phenotyping methods, traits studied, and dependence of GWAS associations on statistical approaches and experimental settings (Table S6-S9; Supplemental Results and Discussion). Several of the most promising gene candidates we found, organized by biological function of orthologs in Arabidopsis, are briefly discussed below.

#### Regulation of cell wall adhesion

Potri.006G276200 encodes a member of the FASCICLIN-LIKE ARABINOGALACTAN (FLA) PROTEIN family and is implicated by a QTL three bases upstream of the transcription start site. We report this association from GMMAT of callus area at week two, the trait with greatest trait with greatest *h^2^_SNP_* as estimated by GEMMA. The significance of this QTL passes the most stringent multiple testing correction method applied – the Bonferroni threshold (α = 0.05) with each individual SNP considered an independent test. No other QTLs associated with this trait meet the same threshold, nor do any other QTLs from GMMAT with any trait in our study.

Many genes in the *FLA* family are differentially expressed during embryogenesis (Costa *et al*., 2019), but their regulation in the context of *in vitro* regeneration has received little study. In Arabidopsis, *FLA1* was found to be upregulated during incubation on callus induction media, while *FLA2* upregulation occurred upon transfer of explants to shoot induction media (Johnson *et al*., 2003). Loss-of-function of *FLA1* was reported to confer an ability for efficient *in vitro* shoot regeneration to the otherwise recalcitrant Col-0 ecotype, while contrarily leading to loss of efficient regeneration in the regenerable ecotype WS (Johnson *et al*., 2011). Thus, effects of differential expression, as is likely to be a consequence of the polymorphism from the SNP location, may be genotype-dependent in poplar as well.

We found an association of shoot development (week four area and PC1) with a window of SNPs including a portion of the promoter and first exon of Potri.019G101900 that is related to Arabidopsis *EXPANSIN B3*. Expansins facilitate the process of pH-dependent cell wall loosening, with various expansins expressed during different stages of development. Perturbations of this gene superfamily have been studied in several plant species, including Arabidopsis, tomatoes, rice, soybean, and tobacco. Overexpression typically produces phenotypes of enhanced growth, such as increased size of plant cells and tissues, as well as reduced fruit firmness. Knockout or knockdown, in contrast, leads to reduced growth and increased firmness (Marowa *et al*., 2016). Expansins are believed to be key regulators of cell wall expansion downstream of auxin, a key hormone for control of regeneration (Majda & Robert, 2018).

#### Regulators of wound-responsive hormone signaling

Potri.006G254100 is a putative homolog of *EIN3-binding F box protein 1 (EBF1)*. Molecular evidence from Arabidopsis suggests that EBF1 facilitates ubiquitin-mediated degradation of ETHYLENE-INSENSITIVE 3 (EIN3) and EIN3-LIKE 1 (EIL1; Potuschak *et al*., 2003; Shen *et al*., 2016) and that this degradation is prevented when EIN3 and EIL1 are stabilized by ETHYLENE-INSENSITIVE 2 (EIN2; An *et al*., 2010). Arabidopsis loss-of-function mutants of *EIN2* were used to supply cotyledon explant material for an *in vitro* regeneration assay, which revealed an approximate fourfold reduction in shoot regeneration in the mutants. The same assay revealed a roughly threefold increase in shoot regeneration with loss-of-function of *HOOKLESS1 (HLS1)*, a gene encoding a putative N-acetyltransferase with a role downstream of *EIN3* in regulating a range of ethylene-regulated traits including apical hook development and *in vitro* regeneration (Chatfield & Raizada, 2008). Also downstream of *EIN3* is positive and negative regulation of numerous genes across at least eight hormone pathways including ethylene signaling and the related jasmonate (JA) and salicylic acid (SA) signaling pathways among others, suggesting that *EIN3* represents a key modulator of hormone crosstalk (Chang *et al*., 2013). Furthermore, *WOUND-INDUCED*

*DEDIFFERENTIATON (WIND)* transcription factors have been shown to regulate ethylene, JA and SA pathways (Iwase *et al*., 2021) and have been identified as regulators of callus and shoot regeneration (Iwase *et al*., 2011, 2016). In support of a key role of *EIN3* in these pathways and in regeneration, we present at least eight gene candidates implicated as interacting directly or indirectly with *EIN3* and upstream regulators of EIN3 (Fig. 9). Our results, considered together with mutant studies in Arabidopsis, suggest that these candidates regulate regeneration by mediating crosstalk between ethylene, JA and SA signaling pathways.

**Figure 9.**
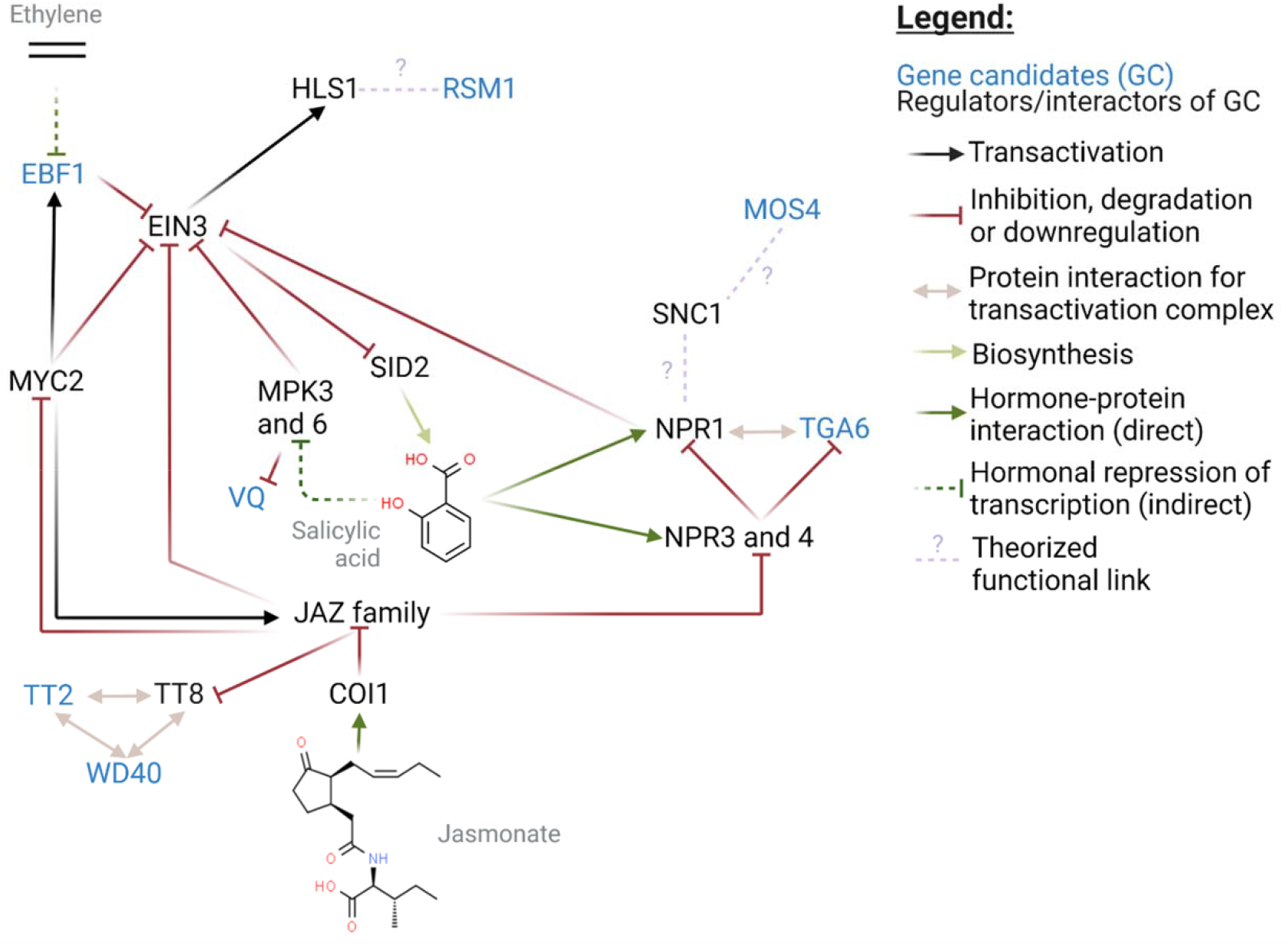
Proposed model integrating roles of *in planta* regeneration gene candidates. Interactions involving Arabidopsis homologs of seven gene candidates (blue) and associated regulators were identified by literature review, providing an understanding of the broader context of hormone crosstalk between ethylene, JA, and SA pathways as they relate to regeneration. Node placement was assisted by the Force Atlas 2 algorithm as implemented in Gephi. Chemical structures for hormones were retrieved from ChemSpider (chemspider.com). Jasmonates are represented by (+)-7-Iso-Jasmonyl-L-isoleucine. This figure was produced using BioRender (biorender.com). Details for each of these gene candidates (blue) can be found in Table 1 and Discussion. Standard acronyms and abbreviations can be found on The Arabidopsis Information Resource (Berardini *et al*., 2015) and are listed in Table S9. Evidence for interactions is summarized in Table S10.

Our GWAS results suggest a central role for SA and genes involved in SA signaling. *NPR1* is a regulator of SA signaling via a mechanism that depends on at least three genes with homologs implicated by QTLs in our GWAS (Fig. 9). Gene candidate Potri.003G194600 encodes a homolog of TGACGT motif transcription factor *TGA6*. *TGA6* and other members of the TGA family have been reported to interact with *NPR1* (Zhang *et al*., 1999; Subramaniam *et al*., 2001; Fan & Dong, 2002; Rochon *et al*., 2006; Boyle *et al*., 2009; Hussain *et al*., 2018) to form a histone acetyltransferase complex responsible for SA-associated epigenetic reprogramming (Jin *et al*., 2018). Simultaneous knockout of functionally redundant *TGA* genes (Zhang *et al*., 2003b) or of *NPR1* (Cao *et al*., 1997) confers a loss of SA signaling, including SA-mediated pathogen resistance. Moreover, a dominant mutation of the putative upstream regulator *SUPPRESSOR OF NPR1, CONSTITUTIVE 1 (SNC1)* confers constitutive SA signaling, dwarf morphology, and enhanced pathogen resistance (Zhang *et al*., 2003a). This phenotype is reversed by loss-of-function of *MODIFIER OF SNC1,4 (MOS4)*, a homolog of our gene candidate Potri.015G041800 (Palma *et al*., 2007).

Additional regulation of *EIN3* is believed to exist via phosphorylation of EIN3 protein by the SA-regulated MAP KINASE 3 (MPK3) and MAP KINASE 6 (MPK6; Yoo *et al*., 2008). MPK3 and MPK6 are also responsible for phosphorylation of many substrates in a VQ MOTIF PROTEIN family related to our gene candidate Potri.002G070600. Although the VQ proteins have not been studied in the context of *in vitro* regeneration to our knowledge, many members are believed to function downstream of pathogen-associated molecular patterns (PAMPs) and upstream of pathogen defense genes (Pecher *et al*., 2014).

Our GWAS results also suggest a central role for anthocyanin and related genes. The SA and JA pathways are linked with anthocyanin signaling by the activity of JAZ proteins in negatively regulating MYB/bHLH/WD40 (MBW) protein complexes responsible for transcriptional regulation of anthocyanin biosynthesis genes (Baudry *et al*., 2004; Qi *et al*., 2011). We report two gene candidates homologous to MBW components, Potri.002G173300 (encoding a WD-40 repeat family protein) and Potri.018G049600 (homolog of *TRANSPARENT TESTA 2*). Although we are unaware of these genes being studied in the context of *in vitro* regeneration, MBWs regulate several steps of anthocyanin biosynthesis downstream of naringenin chalcone, which is produced by CHALCONE SYNTHASE (CHS; Yan *et al*., 2021). CHS knockout in Arabidopsis confers deficient *in vitro* shoot regeneration, with a light-dependent effect. The effects of anthocyanins on shoot regeneration may be mediated by their effects of ROS scavenging (Nameth *et al*., 2013) and/or auxin accumulation (Brown *et al*., 2001).

A functional relationship between *HLS1* (previously described; downstream of *EIN3*) and *RSM1* (homolog of gene candidate Potri.004G155400) has been proposed due to phenotypic similarities between *hls1* and *RSM1*-overexpressing Arabidopsis. Light-deprived seedlings of both mutant lines presented various degrees of reduced hypocotyl length, defective hook formation and defective gravitropism (Hamaguchi *et al*., 2008). However, whereas *HLS1* knockout is known to confer enhanced shoot regeneration in Arabidopsis (Chatfield & Raizada, 2008), the effects of *RSM1* or *RSM* family overexpression or knockout on shoot regeneration have not yet been reported to our knowledge.

Several gene candidates from GWAS appear to affect cytokinin signaling. Potri.010G105600 is a homolog of *ARABIDOPSIS RESPONSE REGULATOR 2 (ARR2)*, which functions shortly downstream of cytokinin. B-type *ARR* genes such as *ARR2* share some degree of functional redundancy and may each positively regulate *in vitro* regeneration via transcriptional upregulation of key developmental genes such as *WUSCHEL* (*WUS;* Xie *et al*., 2018; reviewed by Nagle *et al*., 2018). An additional level of regulation over *WUS* expression exists via the *FANTASTIC FOUR (FAF)* gene family. Overexpression of any of the four *FAF* genes (including *FAF2*, homolog of gene candidate Potri.002G053400) leads to arrest of SAM development, possibly by inhibiting *WUS* expression via an interaction with the feedback loop of regulation between *WUS* and the *WUS* inhibitor *CLUVATA3* (Wahl *et al*., 2010). Shoot meristem development is also regulated by the *CYTOKININ RESPONSE FACTOR (CRF)* gene family (featuring *CRF4*, a homolog of candidate Potri.012G032900), as shown by increased or reduced rosette growth when other members of the *CRF* family are knocked out or overexpressed, respectively. However, these experiments did not feature mutant analysis of the closely related *CRF4* (Raines *et al*., 2016).

#### Reactive oxygen species (ROS) signaling

At least two genes among our candidates appear to have roles in ROS regulation, which may affect regeneration and other developmental processes by mediating post-translational modifications of proteins involved in hormone signaling and/or by affecting levels of oxidative damage to developing tissues. Potri.011G031100 and Potr.012G070400 encode a putative thioredoxin-like protein and a peroxiredoxin, respectively. Although we did not find reports of mutant phenotypes for closely related genes in Arabidopsis in the context of regeneration or related processes, the thioredoxin *DCC1* has been reported to affect *in vitro* shoot regeneration capacity in mutant lines as well as across natural ecotypes of Arabidopsis (Zhang *et al*., 2018b).

## Conclusions

We report a GWAS of *in planta* regeneration in *P. trichocarpa* using a novel system for phenotyping regeneration with computer vision, along with four complementary statistical methods for association mapping. These analyses revealed over 200 candidate genes, strongly implicating regulators of cell adhesion and stress signaling. Further research using gene-editing or other transgenic technologies can validate these associations and investigate the potential of these gene candidates to enhance regeneration in genotypes of poplar and other plants that are recalcitrant to efficient regeneration. While canonical regulators of *in vitro* regeneration tend to be involved in auxin and cytokinin signaling pathways, our results suggest that stress pathways downstream of ethylene, salicylic acid, and jasmonates may be of greatest importance to the mode of *in planta* regeneration that we studied in *P. trichocarpa*. These pathways have received little attention in studies where developmental regulator genes are used to promote regeneration, and would appear to be promising avenues to pursue. Furthermore, at least seven top candidates are likely closely-linked members of a genetic regulatory network, separated from one another by no more than four degrees of direct interactions. This, considered along with the complex nature of *in vitro* regeneration traits, suggests that emerging multi-locus methods and epistasis tests may provide significantly greater insights into the polygenic control of these traits. The scale and precision of this study would not have been possible without computer vision, and our success in identifying established regulators of regeneration demonstrates that semantic segmentation using deep convolutional neural networks shows promise in supporting characterization of regeneration and other complex developmental plant traits.

## Supporting information

Supplemental Methods; Supplemental Results and Discussion

## Acknowledgements

We thank the National Science Foundation Plant Genome Research Program for support (IOS #1546900, Analysis of genes affecting plant regeneration and transformation in poplar), and members of GREAT TREES Research Cooperative at OSU for its support of the Strauss laboratory.

Support for the Poplar GWAS dataset is provided by the U.S. Department of Energy, Office of Science Biological and Environmental Research (BER) via the Center for Bioenergy Innovation (CBI) under Contract No. DE-PS02-06ER64304. The Poplar GWAS Project used resources of the Oak Ridge Leadership Computing Facility and the Compute and Data Environment for Science at Oak Ridge National Laboratory, which is supported by the Office of Science of the U.S. Department of Energy under Contract No. DE-AC05-00OR22725.

This work used the COMET high-performance cluster at the San Diego Supercomputing Center (University of California, San Diego) made available through the Extreme Science and Engineering Discovery Environment (XSEDE), which is supported by National Science Foundation grant number ACI-1548562.

## Contributions of authors

Strauss, Fuxin, Jiang, and Muchero designed and directed the overall study, and obtained funding for its execution; Ma and Peremyslova designed and/or executed the phenotypic analyses; Nagle, Jiang, Yuan, and Kaur created, adapted, and executed the machine vision, computation, and data analysis pipelines; Nagle investigated gene candidates. Niño de Rivera assisted with inspecting results in IGV; Jawdy, Chen, Feng, Yates, Tuskan and Muchero supported acquisition and preparation of the GWAS population and SNP set; Nagle wrote the manuscript with editing from Strauss, and all others contributed further edits and revisions.

## Notes

### Competing Interest Statement

The authors have declared no competing interest.

### Summary of Updates

Details on methods have been added. Methods for SNP set preparation are now in the manuscript. Some additional statistical analysis and interrogation of gene candidates was also conducted.

