## Supplemental Tables 1-5 for "GWAS identifies candidate regulators of in planta regeneration in Populus trichocarpa"

**Supplementary Table 1. Effects of transformations on statistical distributions of *in planta* regeneration traits obtained directly by computer vision.** Raw traits obtained by parsing machine vision outputs into vectors with value of continuous trait for each genotype; all traits were transformed first by excluding zero values, and finally by a Box-Cox transformation. CC = correlation coefficient. Red tones indicate weaker (lower) CC values within rows.

| *Trait category* | *Specific trait* | *Pearson CC of trait and normal dist.* | |
|---|---|---|---|
| | | *Before transf.* | *After transf.* |
| Proportion of area labeled by MV as callus | Week 2 | 0.872 | 0.995 |
| | Week 3 | 0.925 | 0.993 |
| | Week 4 | 0.948 | 0.995 |
| | Week 5 | 0.949 | 0.996 |
| | Growth from week 2-5 | 0.968 | 0.999 |
| Proportion of area labeled by MV as shoot | Week 2 | 0.329 | 0.992 |
| | Week 3 | 0.561 | 0.992 |
| | Week 4 | 0.693 | 0.987 |
| | Week 5 | 0.747 | 0.979 |
| | Growth from week 2-5 | 0.758 | 0.980 |

**Supplementary Table 2. Effects of transformations on statistical distributions of *in planta* regeneration principal component traits obtained by PCA over computer vision traits.** Principal component traits obtained by performing principal component analysis (PCA) to reduce large numbers of raw traits into a smaller number of variables. These traits were processed by a range of transformations, labeled in the "Transf. type" column. Transformation "A" simply involved excluding genotypes which have a zero value for all raw traits, followed by a Box-Cox transformation. Transformation "B" additionally featured removal of outliers prior to Box-Cox, while "C" involved thresholding at an inflection point (elbow) prior to Box-Cox. CC = correlation coefficient. Red tones indicate weaker (lower) CC values within rows.

| *Traits reduced by PCA* | *PC #* | *Transf. type* | *Pearson CC of trait and normal dist.* | |
|---|---|---|---|---|
| | | | *Before transf.* | *After transf.* |
| Four callus traits; Area labeled as callus at all four timepoints | PC1 | A | 0.955 | 0.995 |
| | PC2 | B | 0.965 | 0.992 |
| | PC3 | B | 0.966 | 0.996 |
| | PC4 | B | 0.958 | 0.996 |
| Eight callus/shoot traits; Area labeled as callus or shoot at all four timepoints | PC1 | A | 0.940 | 0.992 |
| | PC2 | C | 0.958 | 0.990 |
| | PC3 | B | 0.982 | 0.993 |
| | PC4 | B | 0.945 | 0.995 |
| Four shoot traits; Area labeled as shoot at all four timepoints | PC1 | A | 0.740 | 0.986 |
| | PC2 | B | 0.797 | 0.963 |
| | PC3 | B | 0.794 | 0.974 |
| | PC4 | B | 0.448 | 0.978 |

**Supplementary Table 3. Relationships between *in planta* regeneration traits, geography and theoretical ancestral subpopulation.** Shown are *p*-values obtained by regression of Box-Cox-transformed traits over Q matrix (fastSTRUCTURE with K = 7 model) and latitude of origin for each clone. Red tones indicate more significant (lower) *p*–values. A single *p*-value is listed for the intercept and effects for each of seven subpopulations because the *p*-values for the intercept and subpopulation effects were the same for each given trait. The directions of relationships between each trait and latitude can be seen in Fig. S8.

| Trait | *p*-values for fixed effects | |
|---|---|---|
| | Intercept / Subpopulations | Latitude |
| Callus Area (wk. 2) | 0.1144 | 0.0162 |
| Callus Area (wk. 3) | 0.0291 | 0.0514 |
| Callus Area (wk. 4) | 0.0121 | 0.0039 |
| Callus Area (wk. 5) | 0.3443 | 0.0409 |
| Callus Area (growth wk. 2-5) | 0.7257 | 0.1066 |
| Shoot Area (wk. 2) | 0.3575 | 0.7122 |
| Shoot Area (wk. 3) | 0.9049 | 0.3317 |
| Shoot Area (wk. 4) | 0.9317 | 0.3464 |
| Shoot Area (wk. 5) | 0.7997 | 0.4817 |
| Shoot Area (growth wk. 2-5) | 0.8044 | 0.4545 |

**Supplementary Table 4. Narrow-sense SNP heritability of *in planta* regeneration traits as computed by GEMMA.** Traits are listed with their and corresponding heritabilities and transformations. Several traits were analyzed with multiple transformations and/or pre-treatments due to a lack in clarity on which transformations were optimal.

| Trait | Heritability | | Treatment of trait data before GWAS | | | |
|---|---|---|---|---|---|---|
| | $h^2_{SNP}$ | SE($h^2_{SNP}$) | Threshold | Removal of duplicate values | Removal of outliers | Transformation |
| Callus (2w) | 0.469 | 0.124 | None | N | N | Box-Cox |
| Callus and shoot PC1 | 0.418 | 0.102 | None | Y | N | RB-INV |
| Callus and shoot PC1 | 0.398 | 0.099 | None | Y | N | Box-Cox |
| Shoot PC2 | 0.389 | 0.255 | None | Y | Y | Box-Cox |
| Callus (3w) | 0.372 | 0.098 | None | N | N | Box-Cox |
| Callus PC1 | 0.352 | 0.103 | None | Y | N | Box-Cox |
| Shoot PC1 | 0.346 | 0.211 | None | Y | N | Box-Cox |
| Callus PC1 | 0.340 | 0.102 | None | Y | N | RB-INV |
| Shoot (2w) | 0.334 | 0.500 | None | N | N | Box-Cox |
| Shoot PC1 | 0.319 | 0.210 | None | Y | N | RB-INV |
| Callus (4w) | 0.308 | 0.105 | None | N | N | Box-Cox |
| Callus and shoot PC2 | 0.241 | 0.095 | -0.198 | Y | Y | Box-Cox |
| Shoot PC2 | 0.208 | 0.162 | None | Y | N | RB-INV |
| Callus (5w) | 0.179 | 0.076 | None | N | N | Box-Cox |
| Shoot PC2 | 0.176 | 0.280 | -0.0146 | Y | Y | Box-Cox |
| Shoot (4w) | 0.121 | 0.167 | None | N | N | Box-Cox |
| Callus growth (2w-5w) | 0.110 | 0.073 | None | N | N | Box-Cox |
| Callus PC2 | 0.102 | 0.110 | None | Y | N | RB-INV |
| Shoot (3w) | 0.097 | 0.325 | None | N | N | Box-Cox |
| Callus and shoot PC2 | 0.097 | 0.061 | None | Y | N | RB-INV |
| Shoot growth (2w-5w) | 0.080 | 0.140 | None | N | N | Box-Cox |
| Shoot (5w) | 0.054 | 0.119 | None | N | N | Box-Cox |
| Callus PC2 | 0.043 | 0.073 | None | Y | Y | Box-Cox |

**Supplementary Table 5. Tallies and statistics of QTL peaks found significant across methods, traits and significance thresholds.** Summaries of genes encompassing or near QTLs are shown for each combination of GWAS method and trait yielding associations that are significant according to a given criteria (conservative Bonferroni, FDR with $\alpha = 0.10$, or ART-Bonferroni). For SKAT and ART associations, the distance shown is that from SNP window centers to the nearest genes. Otherwise, the distance is that between given SNPs and genes.

| Grouping of QTLs by significance and distance | Trait | Method | N genes | Positions of QTL peaks relative to nearest gene | | | | |
|---|---|---|---|---|---|---|---|---|
| | | | | Avg. distance (bp) | Median distance (bp) | Percent intergenic | Percent upstream | Percent downstream |
| All QTLs passing Bonf. | Callus (wk. 2) | GMMAT | 1 | 3 | 3 | 100% | 100% | 0% |
| | Callus (wk. 5) | GEMMA | 1 | 3,947 | 3,947 | 100% | 0% | 100% |
| | Shoot (wk. 4) | MTMCSKAT | 3 | 8,361 | 2,710 | 67% | 33% | 33% |
| | Shoot PC1 | MTMCSKAT | 1 | 29,248 | 29,248 | 100% | 100% | 0% |
| | Shoot PC2 | GEMMA | 2 | 6,911 | 6,911 | 50% | 50% | 0% |
| | Shoot PC2 | MTMCSKAT | 5 | 7,515 | 5,966 | 100% | 80% | 20% |
| All QTLs passing Bonf. within 5kb of gene | Callus (wk. 2) | GMMAT | 1 | 3 | 3 | 100% | 100% | 0% |
| | Callus (wk. 5) | GEMMA | 1 | 3,947 | 3,947 | 100% | 0% | 100% |
| | Shoot (wk. 4) | MTMCSKAT | 2 | 1,355 | 1,355 | 50% | 0% | 50% |
| | Shoot PC2 | GEMMA | 1 | 0 | 0 | 0% | 0% | 0% |
| | Shoot PC2 | MTMCSKAT | 1 | 14 | 14 | 100% | 0% | 100% |
| All QTLs passing FDR (α = 0.10) and/or Bonf. | Callus (wk. 2) | GMMAT | 1 | 3 | 3 | 100% | 100% | 0% |
| | Callus (wk. 2) | MTMCSKAT | 1 | 0 | 0 | 0% | 0% | 0% |
| | Callus (wk. 3) | FarmCPUpp | 1 | 2,839 | 2,839 | 100% | 100% | 0% |
| | Callus (wk. 5) | GEMMA | 7 | 2,207 | 1,850 | 100% | 57% | 43% |
| | Callus growth (wk. 2-5) | MTMCSKAT | 1 | 0 | 0 | 0% | 0% | 0% |
| | Callus, Shoot PC2 | MTMCSKAT | 1 | 27,546 | 27,546 | 100% | 100% | 0% |
| | Shoot (wk. 4) | MTMCSKAT | 3 | 8,361 | 2,710 | 67% | 33% | 33% |
| | Shoot PC1 | MTMCSKAT | 16 | 7,576 | 946 | 63% | 38% | 25% |
| | Shoot PC2 | GEMMA | 11 | 2,802 | 477 | 73% | 36% | 36% |
| | Shoot PC2 | MTMCSKAT | 5 | 7,515 | 5,966 | 100% | 80% | 20% |
| All QTLs passing FDR (α = 0.10) and/or Bonf. within 5kb of gene | Callus (wk. 3) | FarmCPUpp | 1 | 2,839 | 2,839 | 100% | 100% | 0% |
| | Callus (wk. 5) | GEMMA | 7 | 2,207 | 1,850 | 100% | 57% | 43% |
| | Shoot PC2 | GEMMA | 8 | 657 | 282 | 63% | 25% | 38% |
| | Callus (wk. 2) | GMMAT | 1 | 3 | 3 | 100% | 100% | 0% |
| | Callus (wk. 2) | MTMCSKAT | 1 | 0 | 0 | 0% | 0% | 0% |
| | Callus growth (wk. 2-5) | MTMCSKAT | 1 | 0 | 0 | 0% | 0% | 0% |
| | Shoot (wk. 4) | MTMCSKAT | 2 | 1,355 | 1,355 | 50% | 0% | 50% |
| | Shoot PC1 | MTMCSKAT | 11 | 996 | 0 | 45% | 18% | 27% |
| | Shoot PC2 | MTMCSKAT | 1 | 14 | 14 | 100% | 0% | 100% |
| All QTLs passing ART-Bonf. | Callus (wk. 4) | GMMAT | 13 | 4,064 | 1879 | 0% | 77% | 23% |
| | Callus (wk. 5) | GMMAT | 6 | 1,114 | 1040 | 17% | 50% | 33% |
| | Callus, Shoot PC2 | GMMAT | 6 | 1,669 | 1620 | 17% | 17% | 67% |
| | Callus (wk. 2) | GEMMA | 8 | 3,411 | 3,394 | 100% | 88% | 13% |
| | Callus (wk. 3) | GEMMA | 5 | 1,951 | 1,750 | 80% | 60% | 20% |

| | | | | | | | | |
|---|---|---|---|---|---|---|---|---|
| | Callus (wk. 4) | GEMMA | 8 | 1,868 | 1,522 | 63% | 38% | 25% |
| | Callus growth (wk. 2-5) | GEMMA | 11 | 3,065 | 1,997 | 91% | 73% | 18% |
| | Callus PC1 | GEMMA | 6 | 2,774 | 2,782 | 83% | 50% | 33% |
| | Callus PC2 | GEMMA | 20 | 2,006 | 1,624 | 90% | 55% | 35% |
| | Callus, Shoot PC1 | GEMMA | 3 | 8,135 | 5,222 | 100% | 100% | 0% |
| | Callus, Shoot PC2 | GEMMA | 17 | 1,973 | 1,658 | 88% | 71% | 18% |
| | Shoot (wk. 4) | GEMMA | 6 | 4,630 | 1,308 | 67% | 67% | 0% |
| | Shoot PC1 | GEMMA | 19 | 2,192 | 1,804 | 84% | 58% | 26% |
| | Shoot PC2 | GEMMA | 27 | 7,503 | 1,784 | 81% | 48% | 33% |
| All QTLs passing ART-Bonf. within 5kb of gene | Callus (wk. 4) | GMMAT | 9 | 1,868 | 1427 | 0% | 78% | 22% |
| | Callus (wk. 5) | GMMAT | 6 | 1,114 | 1040 | 17% | 50% | 33% |
| | Callus, Shoot PC2 | GMMAT | 6 | 1,669 | 1620 | 17% | 17% | 67% |
| | Callus (wk. 2) | GEMMA | 5 | 1,851 | 1,059 | 100% | 80% | 20% |
| | Callus (wk. 3) | GEMMA | 4 | 957 | 908 | 75% | 75% | 0% |
| | Callus (wk. 4) | GEMMA | 7 | 1,314 | 1,489 | 57% | 29% | 29% |
| | Callus growth (wk. 2-5) | GEMMA | 9 | 1,886 | 1,620 | 89% | 78% | 11% |
| | Callus PC1 | GEMMA | 5 | 2,232 | 1,741 | 80% | 60% | 20% |
| | Callus PC2 | GEMMA | 19 | 1,716 | 1,348 | 89% | 53% | 37% |
| | Callus, Shoot PC1 | GEMMA | 1 | 2,510 | 2,510 | 100% | 100% | 0% |
| | Callus, Shoot PC2 | GEMMA | 17 | 1,973 | 1,658 | 88% | 71% | 18% |
| | Shoot (wk. 4) | GEMMA | 4 | 654 | 445 | 50% | 50% | 0% |
| | Shoot PC1 | GEMMA | 18 | 1,887 | 1,716 | 83% | 56% | 28% |
| | Shoot PC2 | GEMMA | 23 | 1,556 | 1,151 | 78% | 48% | 30% |
