## Supplemental Table 9 for "GWAS identifies candidate regulators of in planta regeneration in Populus trichocarpa"

**Supplementary Table 9. Abbreviations and acronyms for Arabidopsis homologs of gene candidates and their interactors**

| *Acronym/abbreviation* | *Full name* | *Other acronym/abbreviation* | *Other full name* |
|---|---|---|---|
| HLS1 | HOOKLESS 1 | COP3 | CONSTITUTIVE PHOTOMORPHOGENIC 3 |
| RSM3 | RADIALIS-LIKE SANT/MYB 3 | MEE3 | MATERNAL EFFECT EMBRYO ARREST 3 |
| EBF1 | EIN3-BINDING F BOX PROTEIN 1 | | |
| EIN3 | ETHYLENE-INSENSITIVE3 | | |
| MOS4 | MODIFIER OF SNC1,4 | | |
| MYC2 | MYELOCYTOMATOSIS-LIKE 2 | JAI1 | JASMONATE INSENSITIVE 1 |
| MPK3 | MITOGEN-ACTIVATED PROTEIN KINASE 3 | | |
| MPK6 | MITOGEN-ACTIVATED PROTEIN KINASE 6 | | |
| VQ (family) | VQ motif-containing protein family | | |
| SID1 | SALICYLIC ACID INDUCTION DEFICIENT 1 | EDS5 | ENHANCED DISEASE SUSCEPTIBILITY 5 |
| SNC1 | SUPPRESSOR OF NPR1-1, CONSTITUTIVE 1 | | |
| NPR1 | NONEXPRESSER OF PR GENES 1 | SAI1 | SALICYLIC ACID INSENSITIVE 1 |
| TGA6 | TGACG SEQUENCE-SPECIFIC BINDING PROTEIN 6 | | |
| NPR3 | NONEXPRESSER OF PR GENES 3 | | |
| NPR4 | NONEXPRESSER OF PR GENES 4 | | |
| JAZ (family) | JASMONATE-ZIM-DOMAIN PROTEIN family | | |
| TT8 | TRANSPARENT TESTA 8 | | |
| TT2 | TRANSPARENT TESTA 2 | MYB123 | MYB DOMAIN PROTEIN 123 |
| WD40 (family) | Transducin/WD40 repeat-like protein superfamily | | |
| COI1 | CORONATINE INSENSITIVE1 | | |
