## Supplemental Figures 1-13 for "GWAS identifies candidate regulators of in planta regeneration in Populus trichocarpa"

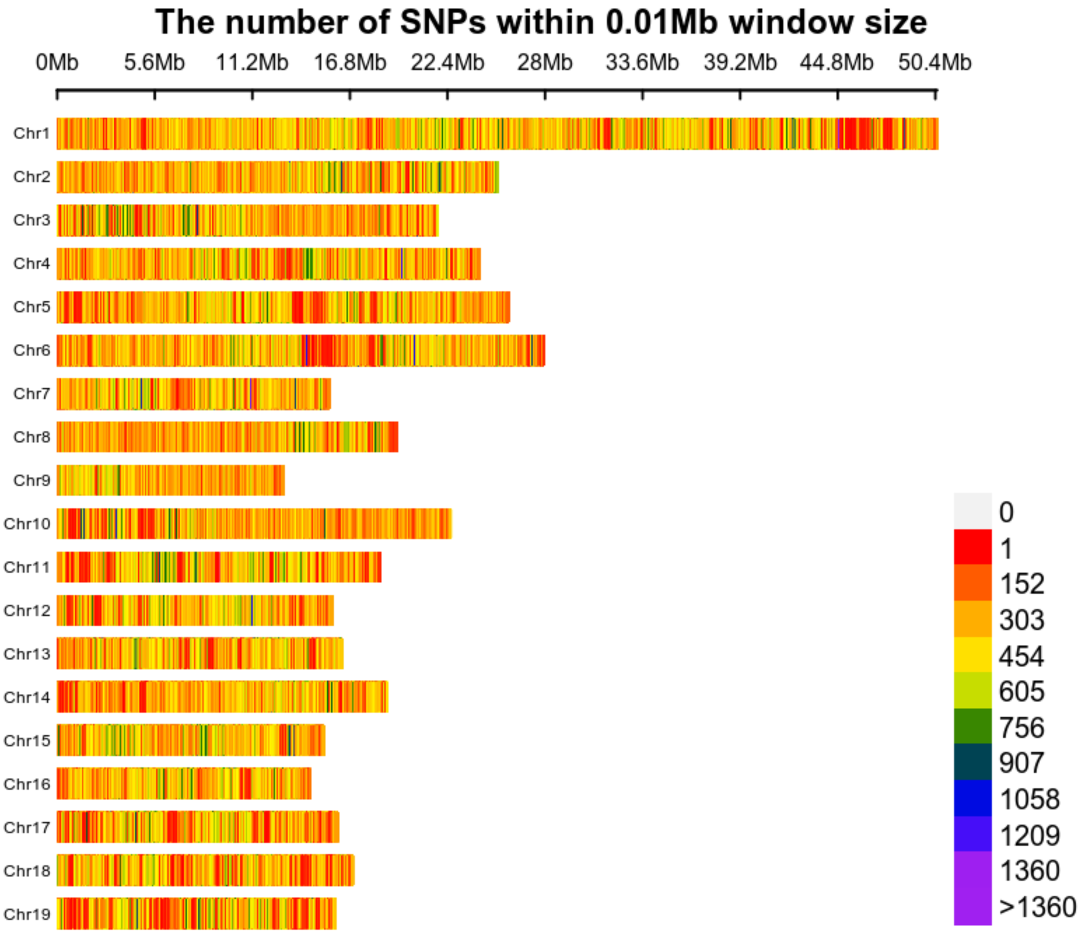


**Supplementary Figure 1. SNP density plot.** The density of SNPs is shown for a SNP set of approximately 10.3 million SNPs, filterted on the basis of MAF (MAF > 0.05), and limited to SNPs on assembled contiguous chromosomes. This plot is shown with a bin size of 10kb and was generated using `CMplot` (R).

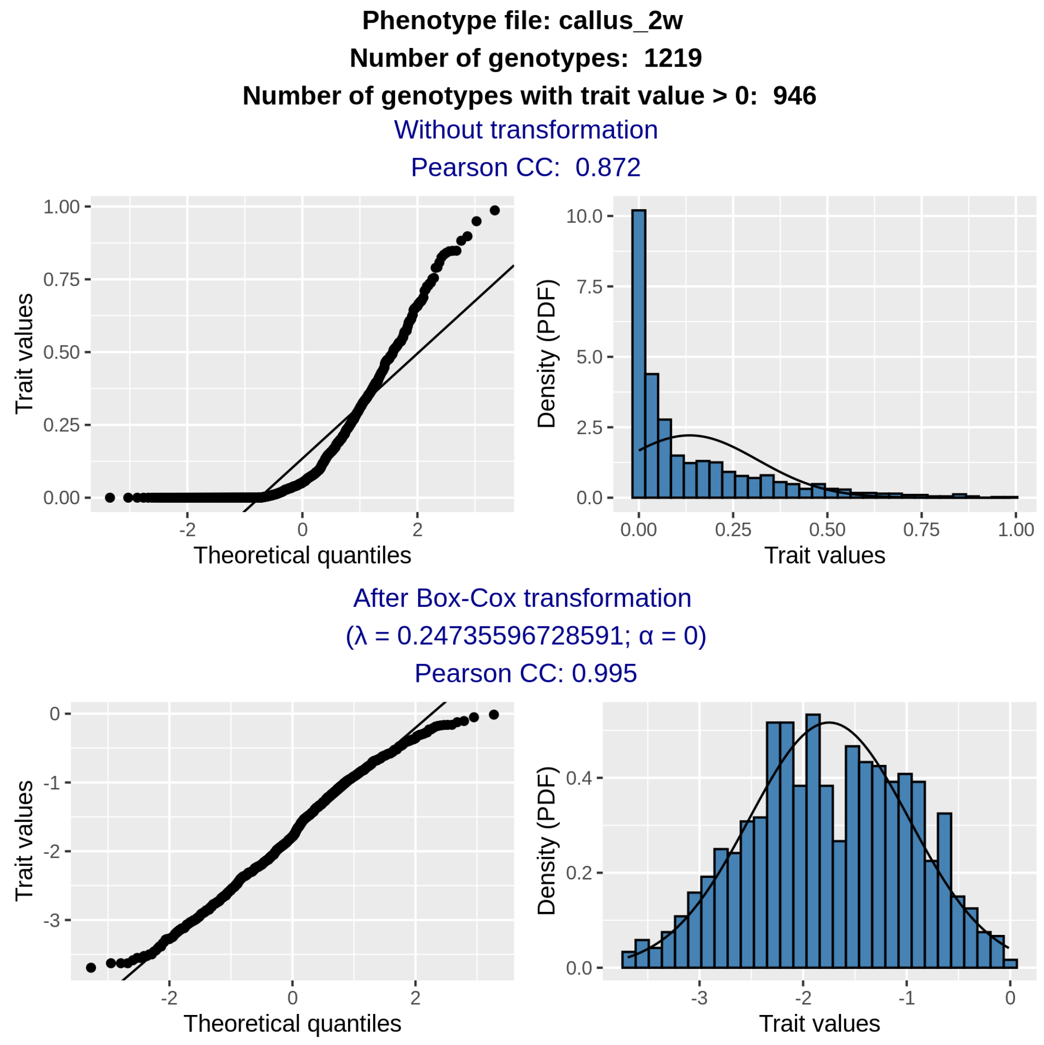


**Supplementary Figure 2. Trait before and after Box-Cox transformation, given for the example of Callus Area at week 2.** Zero values were excluded before (above) and after (below) transformation. The QQ plots (left) show quantiles of the distribution plotted against theoretical quantiles of normal distributions with the same mean and standard deviation. These theoretical normal distributions are also represented by a black line in the histograms (right).

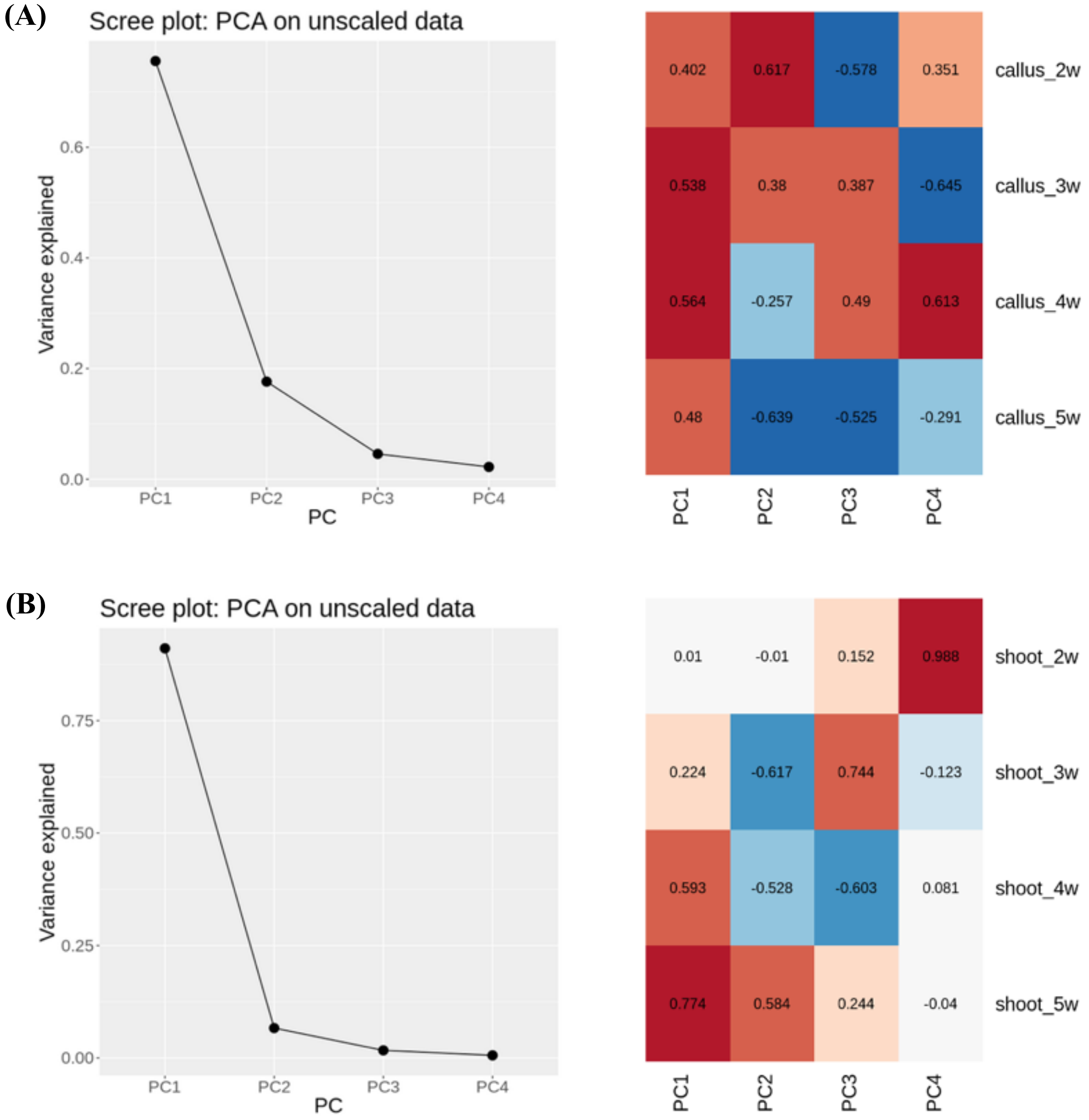


**Supplementary Figure 3. Results from PCA over *in planta* regeneration traits.** Results are shown for two PCA batches: over all callus traits **(A)** and shoot traits **(B)**. Scree plots (left) show most variation is explained by the top two principal components. Heat maps of loadings from PCA (right) show contributions of each trait to each PC.

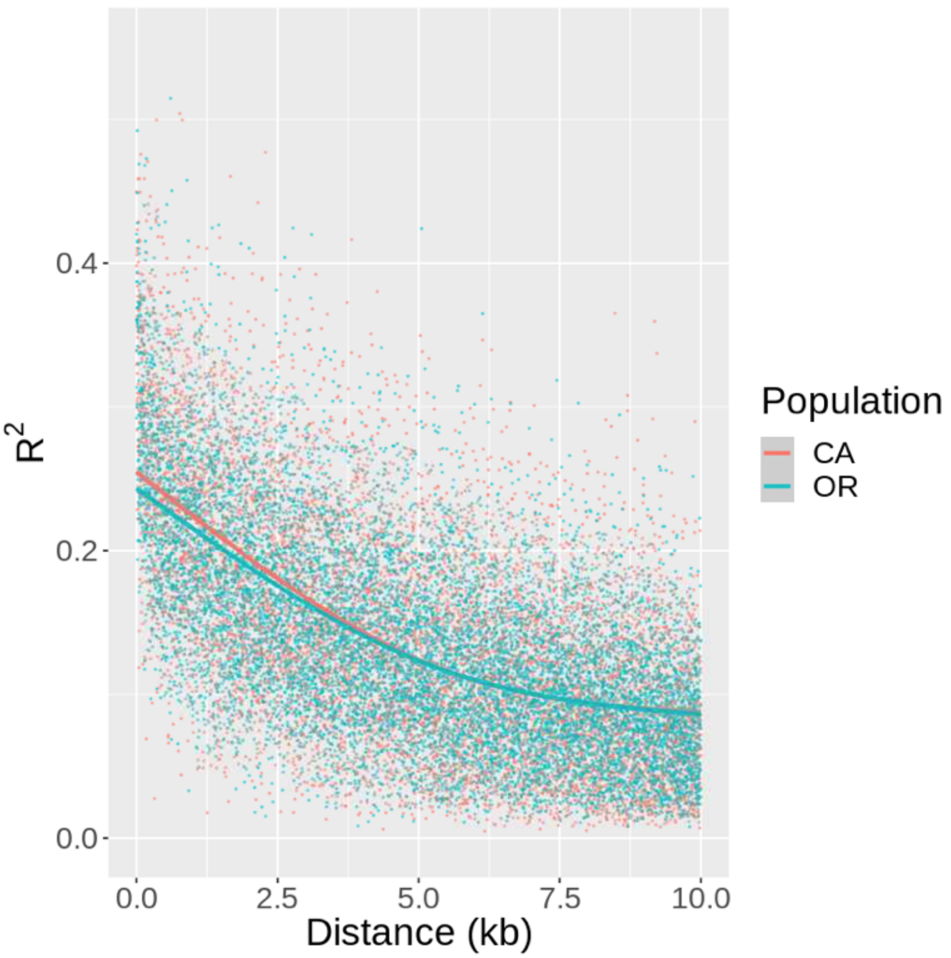


**Supplementary Figure 4. Linkage disequilibrium decay curves.** Graphs are shown for Oregon ("OR") subpopulation (group 5 in Fig. S7) and California ("CA") subpopulation (group 2 in Fig. S7). Primary subpopulations were determined using fastSTRUCTURE, with a K = 7 model (Supplemental Methods). Each point represents the mean $R^2$ between SNPs of a given distance (x-axis). Lines represent LD decay as computed with a spline function. The difference between the CA and OR rates of LD decay was statistically significant ($P < 0.001$) based on 1000 permutations of a spline model (Supplemental Results and Discussion).

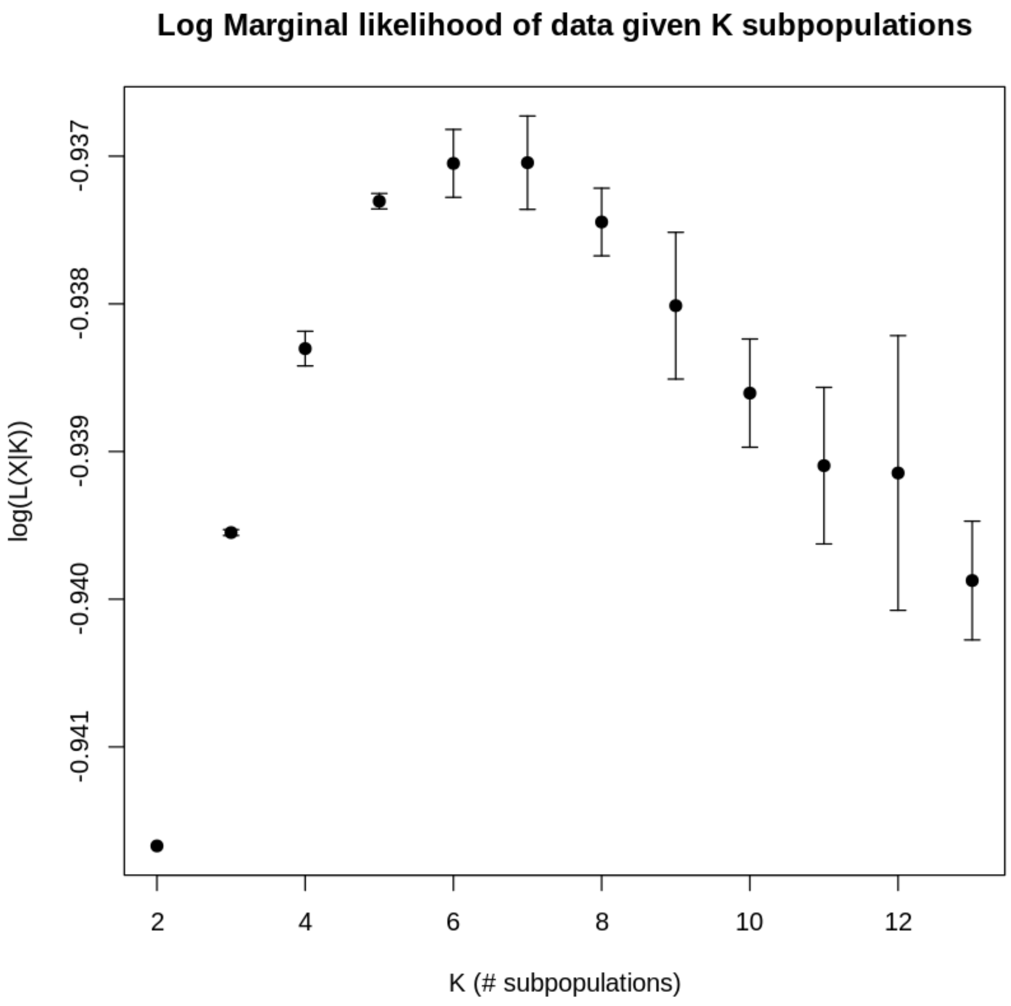


**Supplementary Figure 5. Analysis of subpopulations.** fastSTRUCTURE was performed with 10 replicates, indicating that population structure is best represented with six or seven subpopulations.

(A) (B)

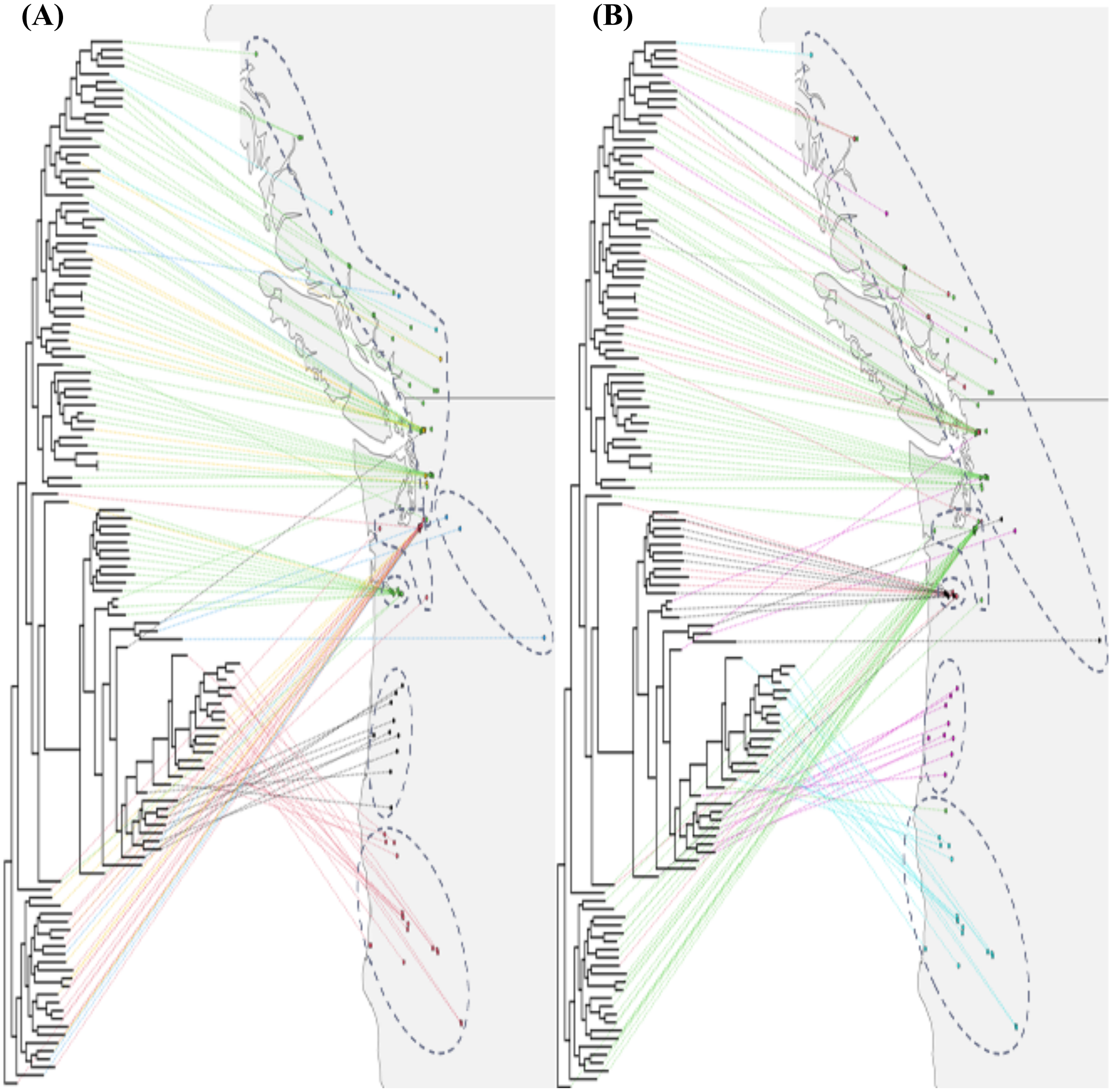

**Supplementary Figure 6. Information on theoretical ancestral subpopulations via fastSTRUCTURE. (A)** K = 7 model and **(B)** K = 6 model. Subpopulations are cross-referenced with both dendrogram and geography, with primary clones colored by primary subpopulation, for 130 randomly selected clones for readability. This plot was produced using the `phytools` R package. For improved readability, dotted demarcations were manually added to indicate approximate groupings based on fastSTRUCTURE clusters and/or phylogenetic groupings.

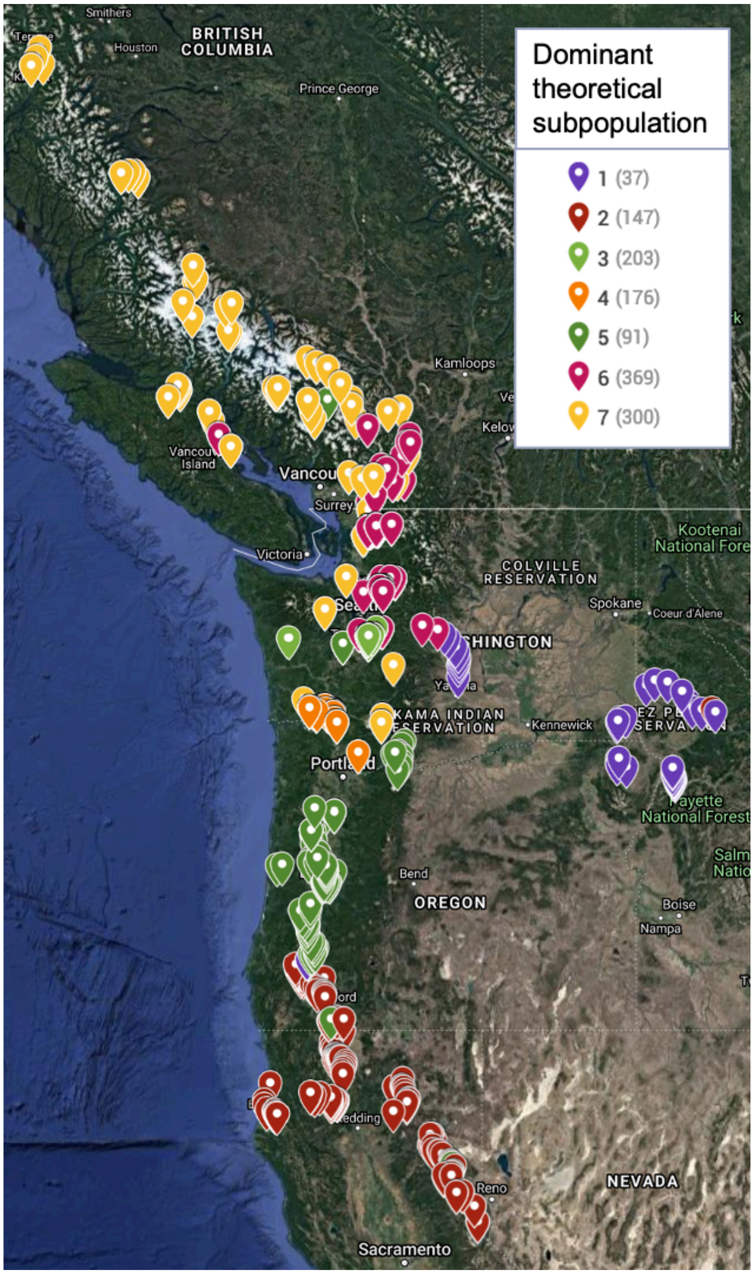


**Supplementary Figure 7. Phylogenetic relationships of natural source populations.** Shown are the theoretical ancestral subpopulations (fastSTRUCTURE, with K=7 model), cross-referenced with geographical locations of the study clones taken from the wild. Data is shown for the 1,301 clones for which location data is available (out of 1,323). Points are labeled by the theoretical subpopulation accounting for the largest portion of ancestry for each clone. The plot was produced by Google Maps (MyMaps).

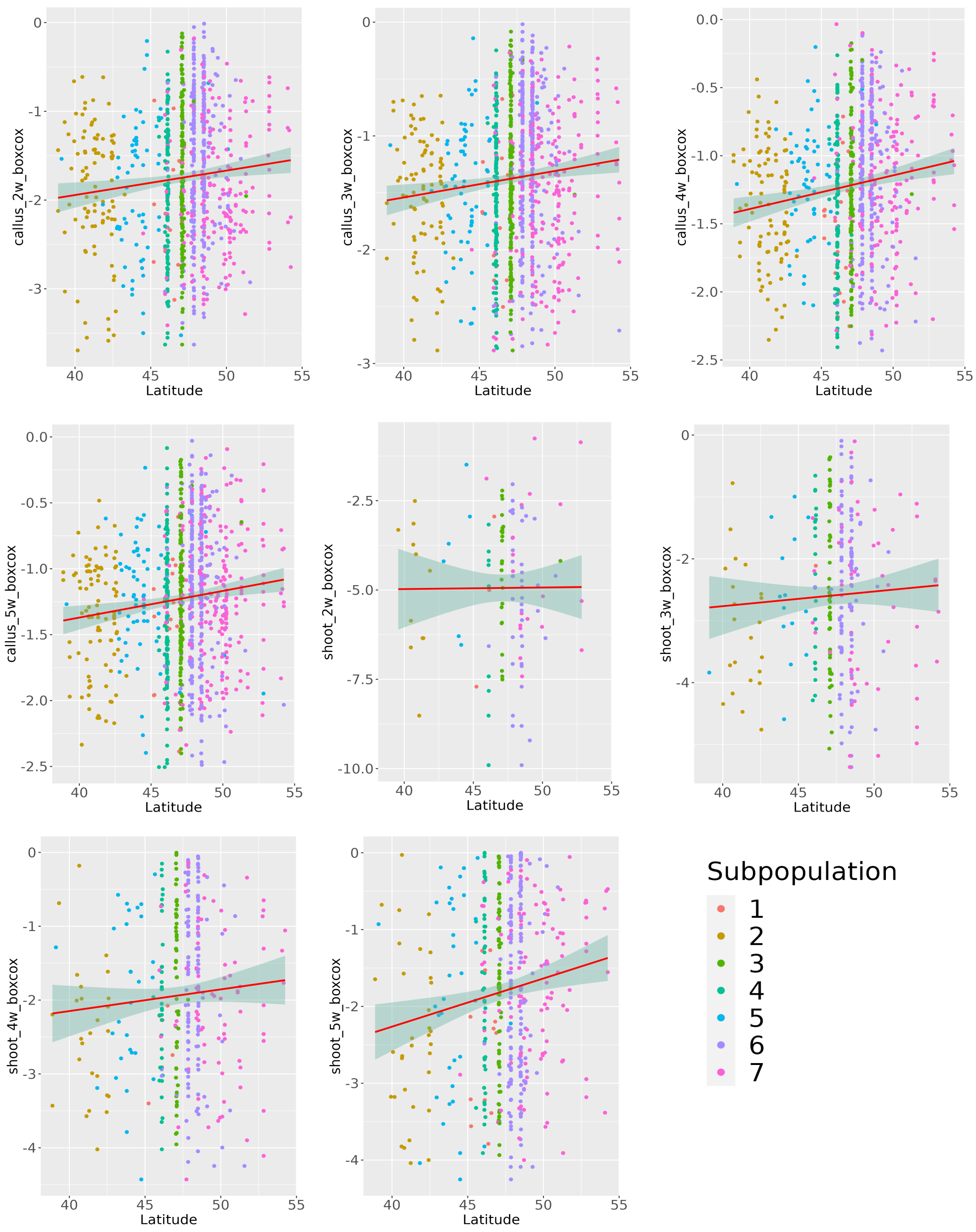


**Supplementary Figure 8. Relationships between callus/shoot area traits and latitude of clone of origin.** Clones are labeled by primary subpopulation (fastSTRUCTURE). A trendline with a 95% confidence interval is displayed for the general relationship between each trait and latitude. These plots were produced using `ggplot2` (R).

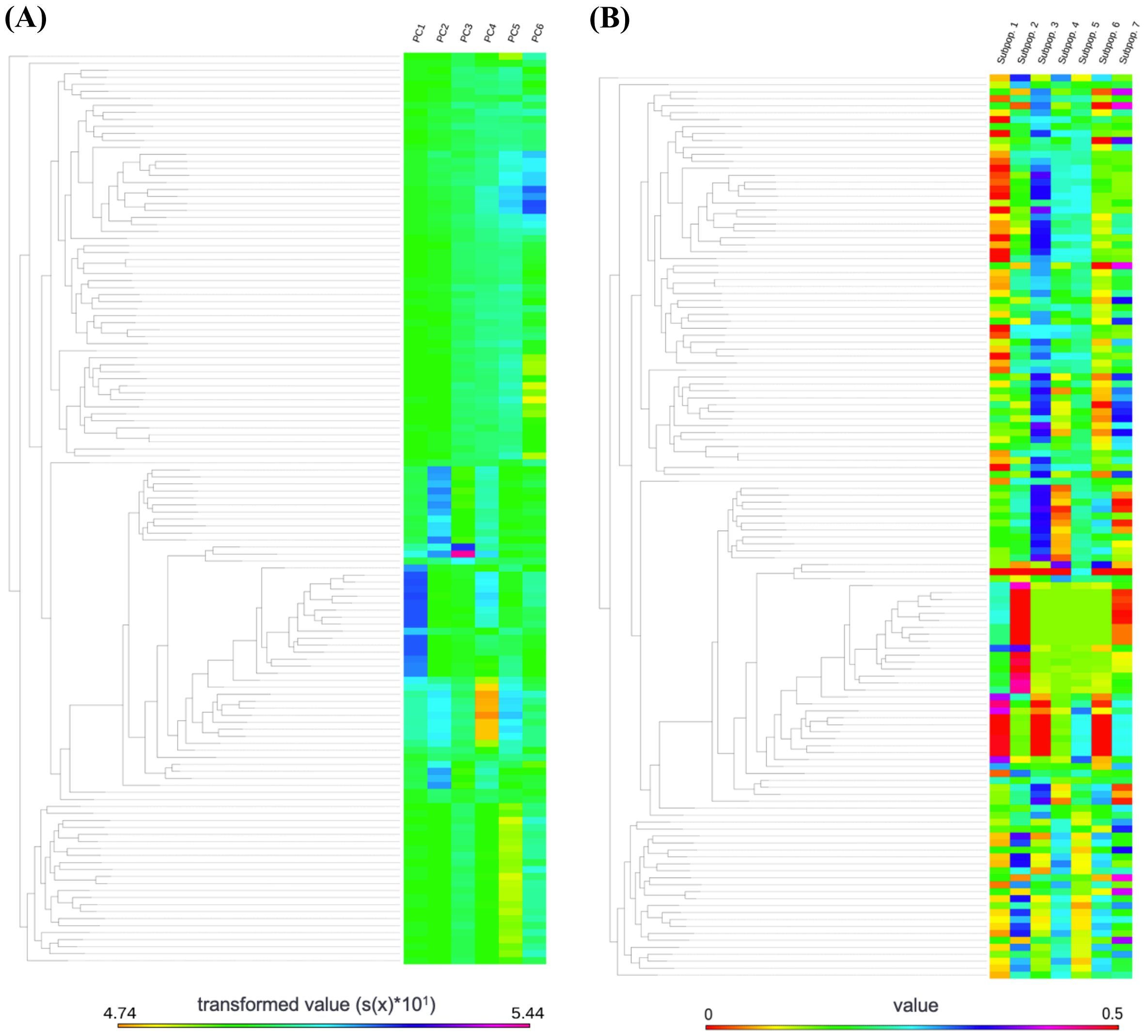


**Supplementary Figure 9. Dendrogram produced by SNPhylo.** Results are cross-referenced with **(A)** top six principal components from PCA over SNP data and **(B)** subpopulations from fastSTRUCTURE (K = 7 model). Data is shown for only 130 randomly selected clones for readability.

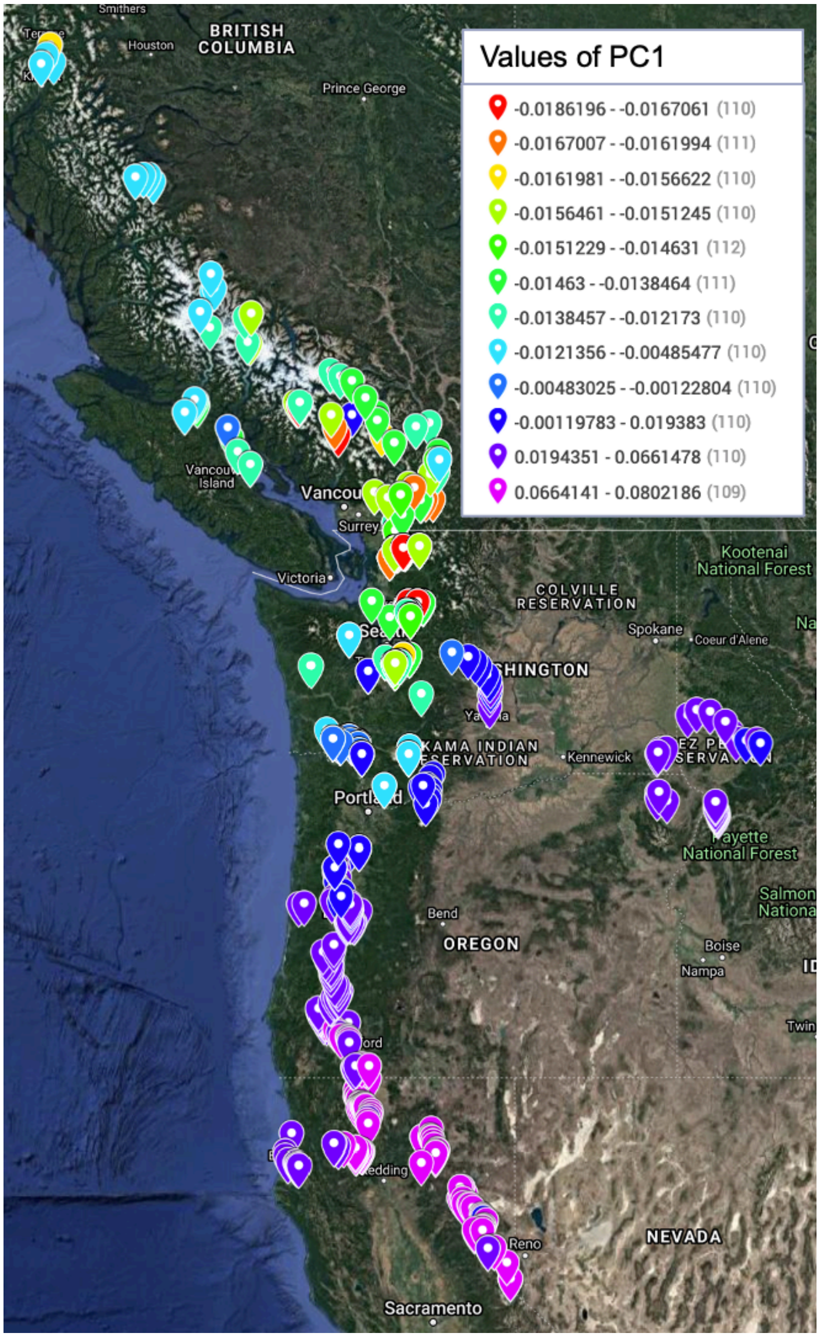


**Supplementary Figure 10. Principal component analysis results cross-referenced with geographical locations of clones.** Data is shown for the 1301 clones for which location data was available (out of 1323). This plot was produced with Google Maps MyMaps.

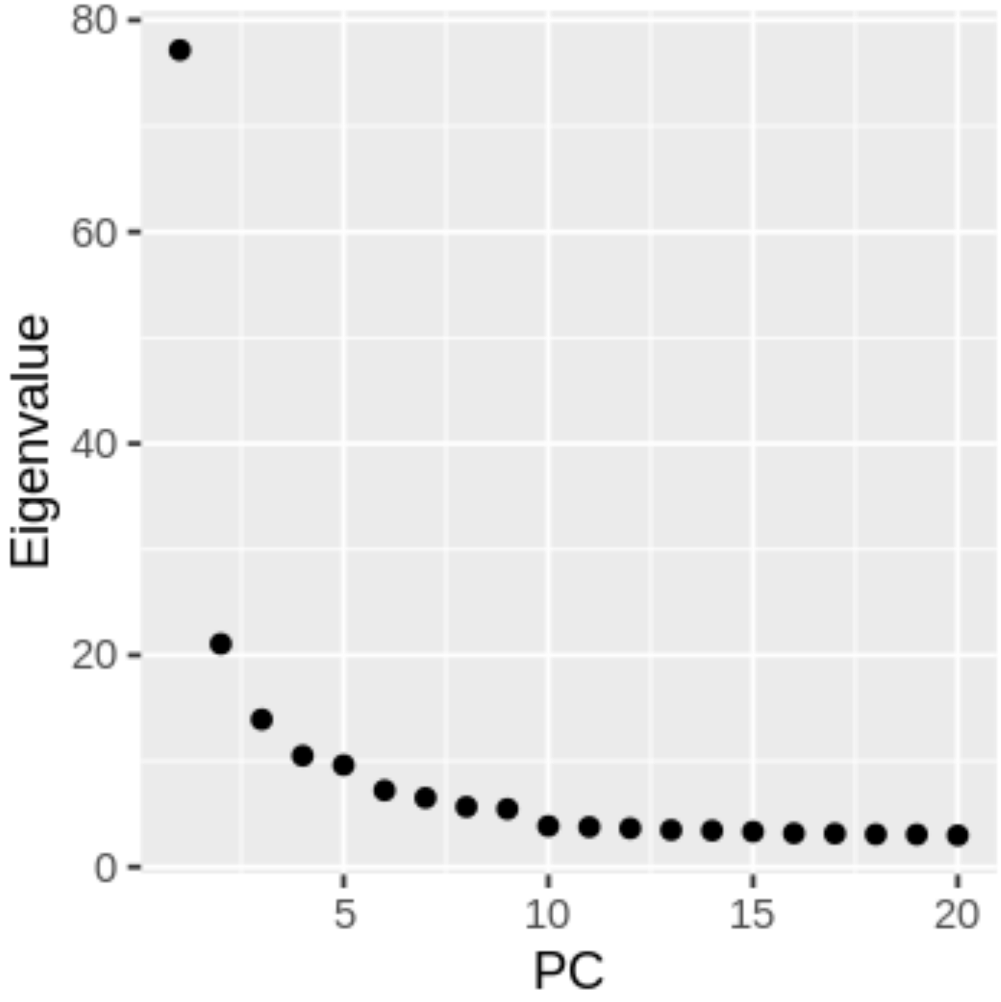


**Supplementary Figure 11. Scree plot from PCA over SNP data.** The scree plot is shown with eigenvalues for each of 20 PCs obtained by PCA over 10.3 M SNPs with MAF > 0.05: PCA was performed using PLINK and the plot was produced with `ggplot2` (R).

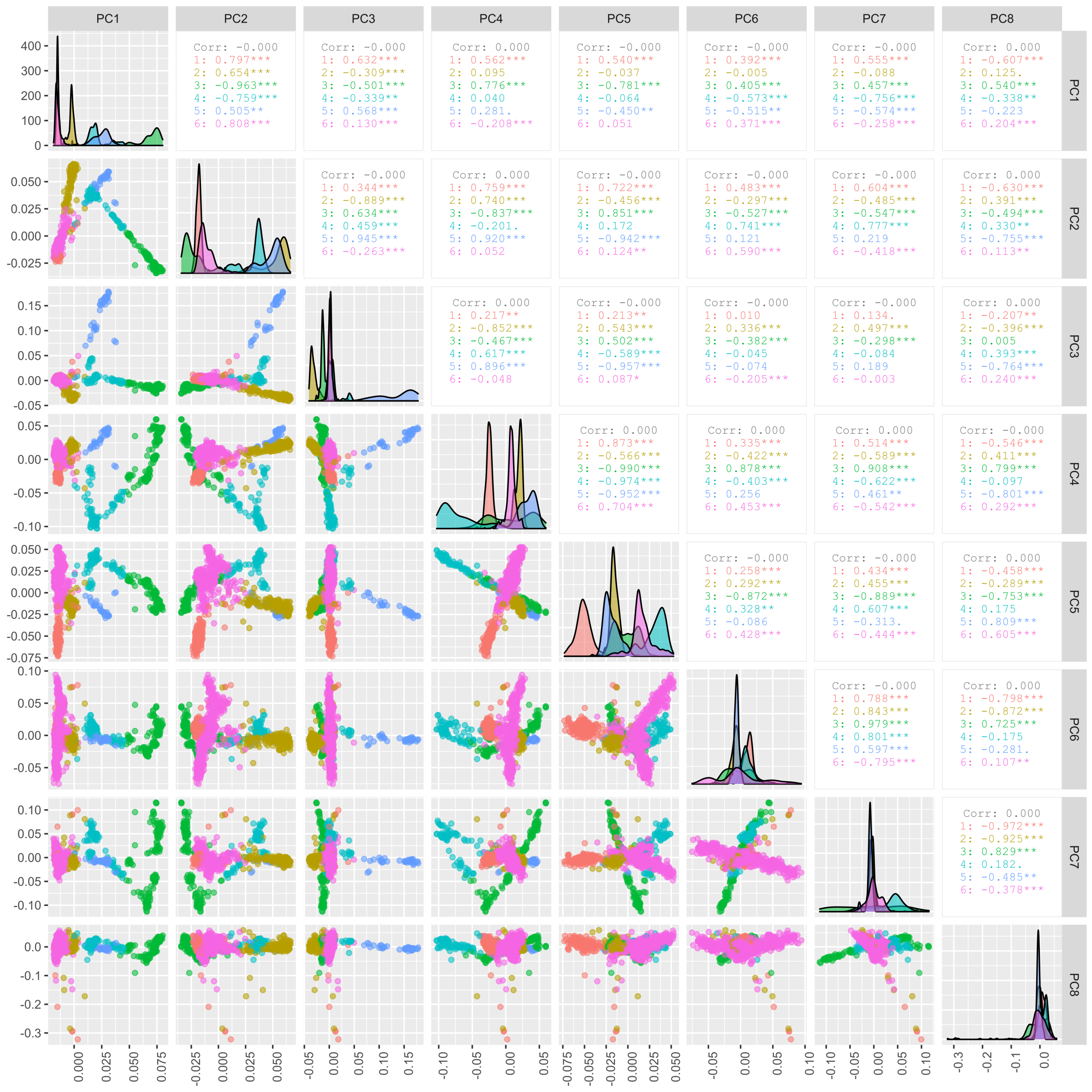


**Supplementary Figure 12. Correlations between PCs from PCA over SNP data.** Correlations are shown between top eight PCs from PCA over 10.3M SNPs with MAF >0.05: K-means clusters (with k = 6) were computed using R and used to group and label samples, with each cluster shown by a different color. This plot was produced with `ggpairs` (R).

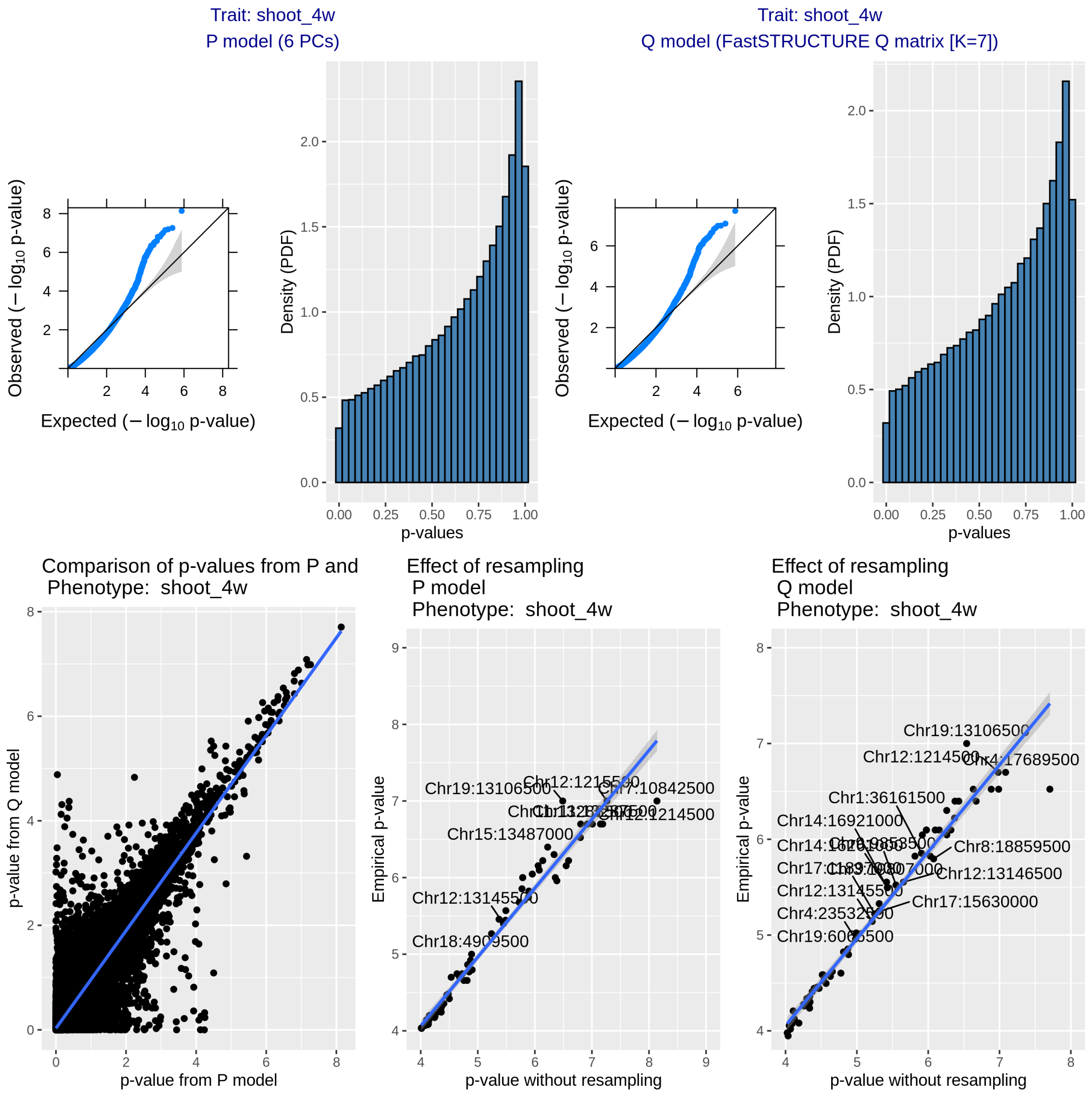


**Supplementary Figure 13. Comparison of P and Q models with SKAT.** Q-Q plots and histograms of *p*-values are shown, along with comparisons of *p*-values between the P and Q models and between empirical *p*-values and *p*-values before resampling for each method
