## Supplemental Figures 14 for "GWAS identifies candidate regulators of in planta regeneration in Populus trichocarpa"

**Supplementary Figure 14. Genome browser views of ART associations.** Association mapping with ART was followed by inspection of peaks and their alignment with gene annotations to find cases where ART peaks implicate specific genes. Nine examples are shown.

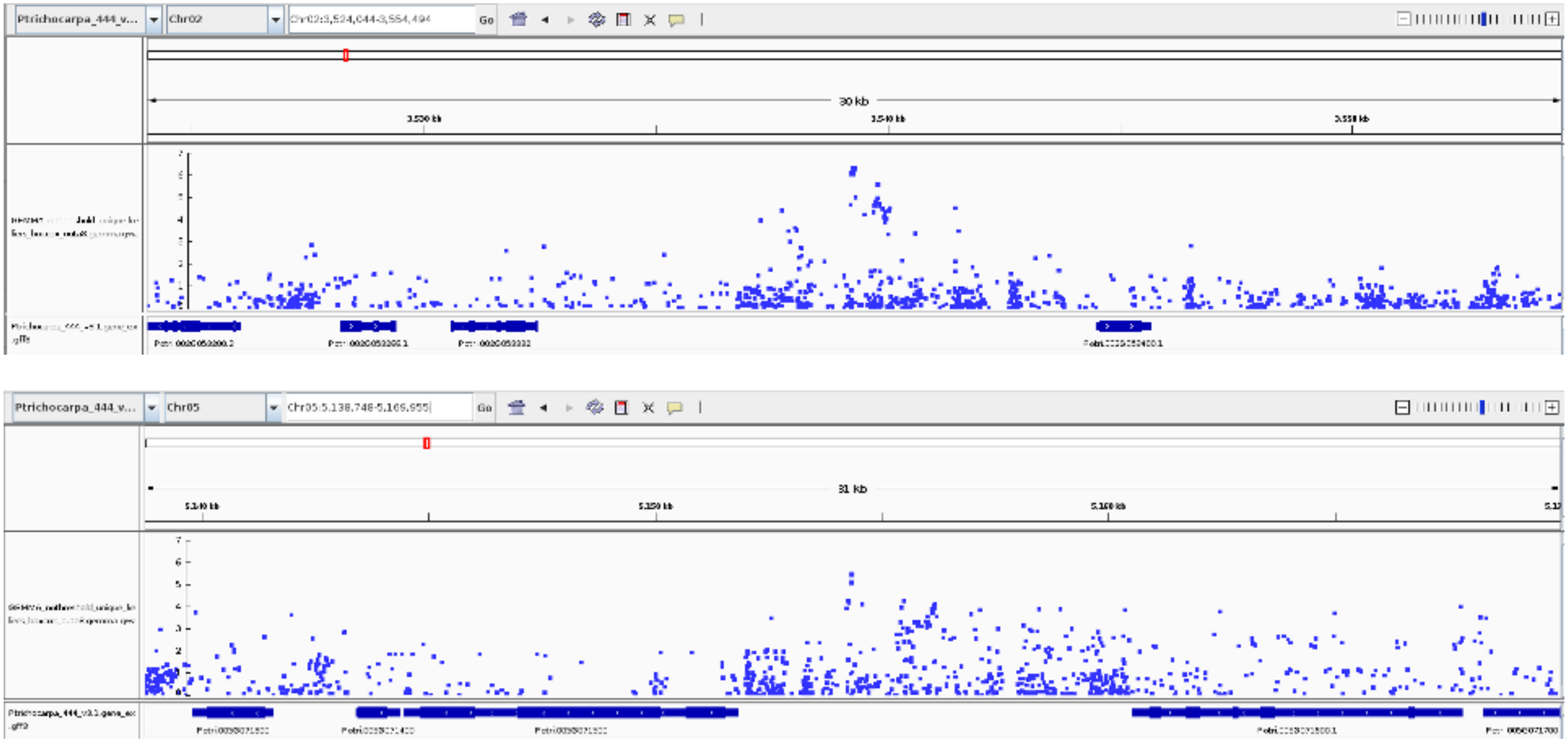

**Supplementary Figure 14. Genome browser views of ART associations (continued)**

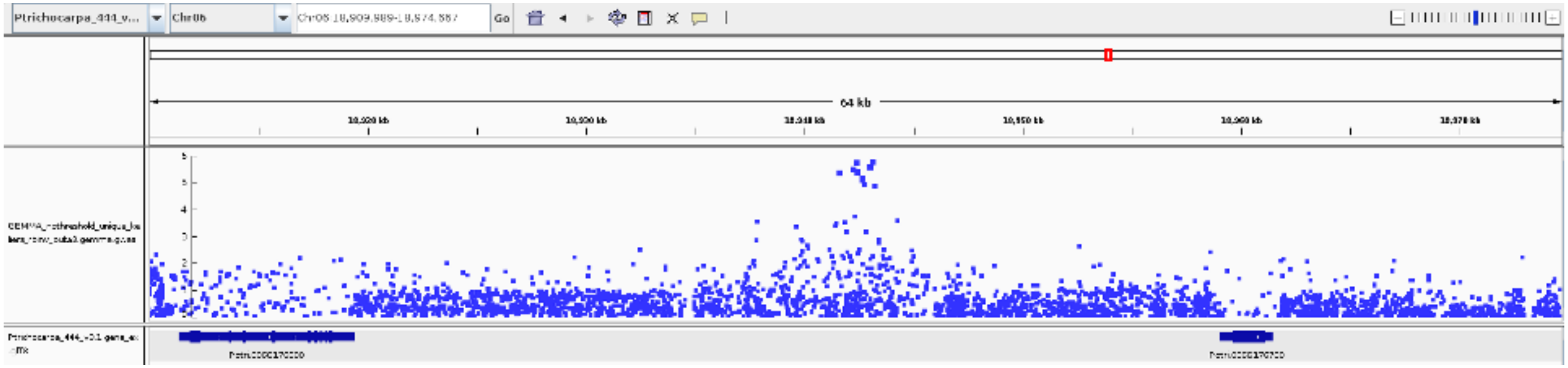
Ptrichocarpa_444_v...
Go
64 kb

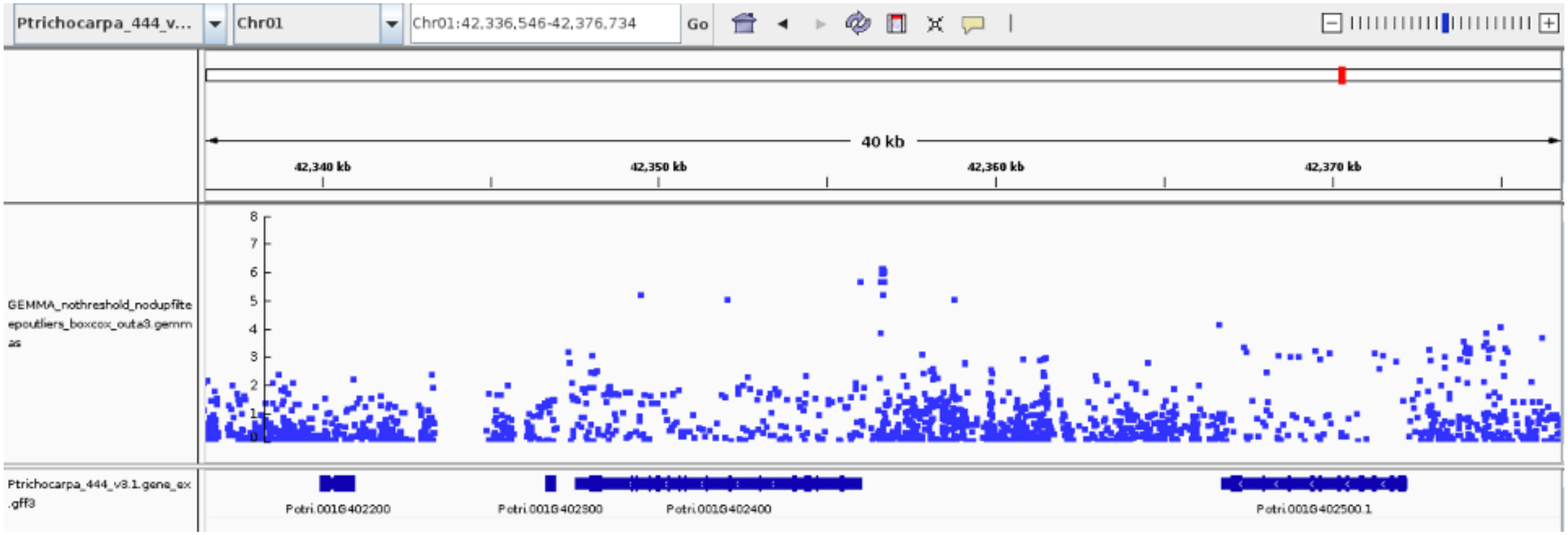
Ptrichocarpa_444_v...
Chr01
Chr01:42,336,546-42,376,734
Go
40 kb
42,340 kb
42,350 kb
42,360 kb
42,370 kb
GEMMA_nothreshold_nodupfilte
epoutliers_boxcox_outa3.gemm
as
8
7
6
5
4
3
2
1
0
Ptrichocarpa_444_v3.1.gene_ex
.gff3
Potri.001G402200
Potri.001G402300
Potri.001G402400
Potri.001G402500.1

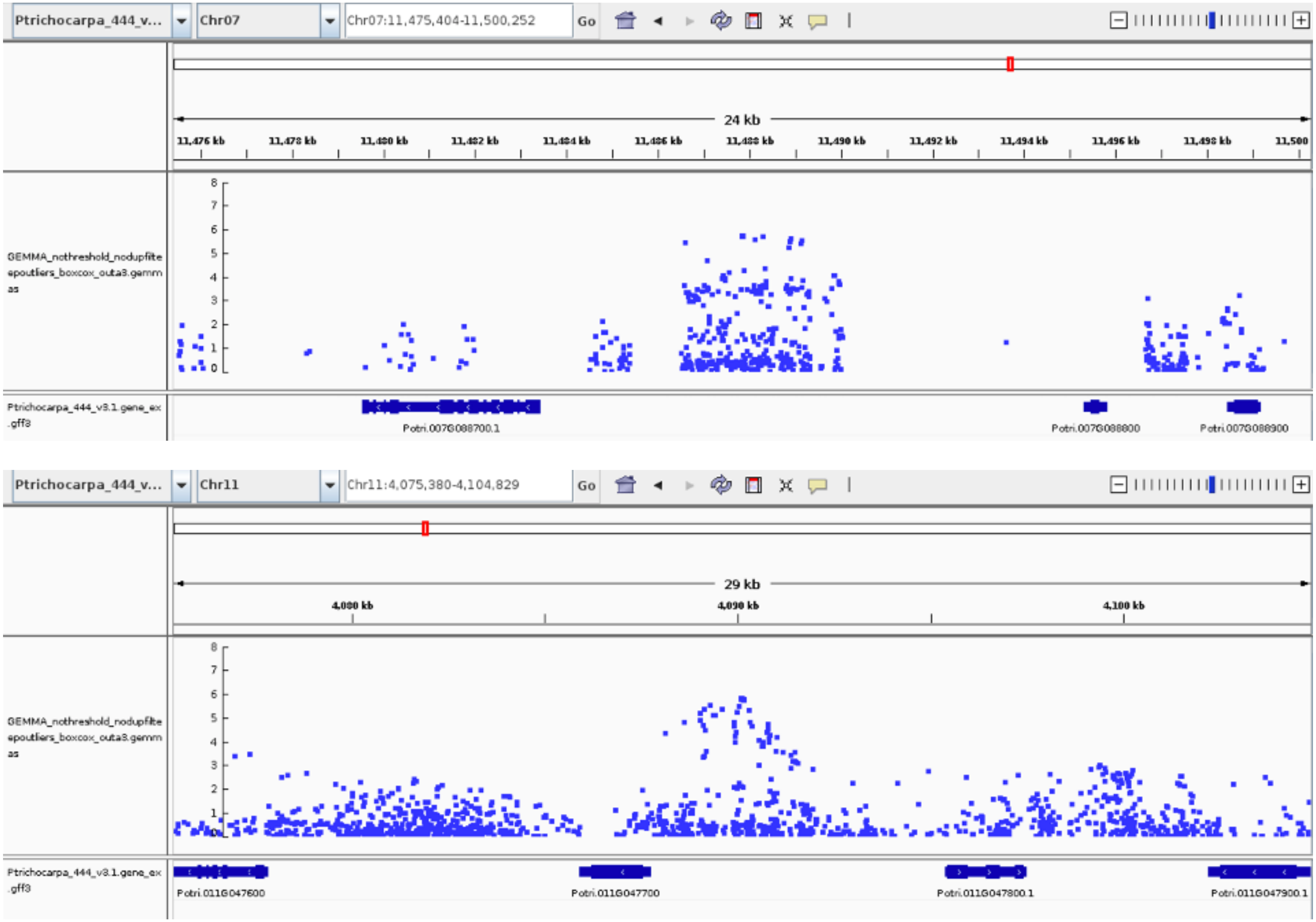

Ptrichocarpa_444_v...
Chr07
Chr07:11,475,404-11,500,252
Go
24 kb
11,476 kb
11,478 kb
11,480 kb
11,482 kb
11,484 kb
11,486 kb
11,488 kb
11,490 kb
11,492 kb
11,494 kb
11,496 kb
11,498 kb
11,500
GEMMA_nothreshold_nodupfilteroutliers_boxcox_outa3.gemmas
Ptrichocarpa_444_v3.1.gene_exons.gff3
Potri.007G088700.1
Potri.007G088800
Potri.007G088900
Ptrichocarpa_444_v...
Chr11
Chr11:4,075,380-4,104,829
Go
29 kb
4,080 kb
4,090 kb
4,100 kb
GEMMA_nothreshold_nodupfilteroutliers_boxcox_outa3.gemmas
Ptrichocarpa_444_v3.1.gene_exons.gff3
Potri.011G047600
Potri.011G047700
Potri.011G047800.1
Potri.011G047900.1

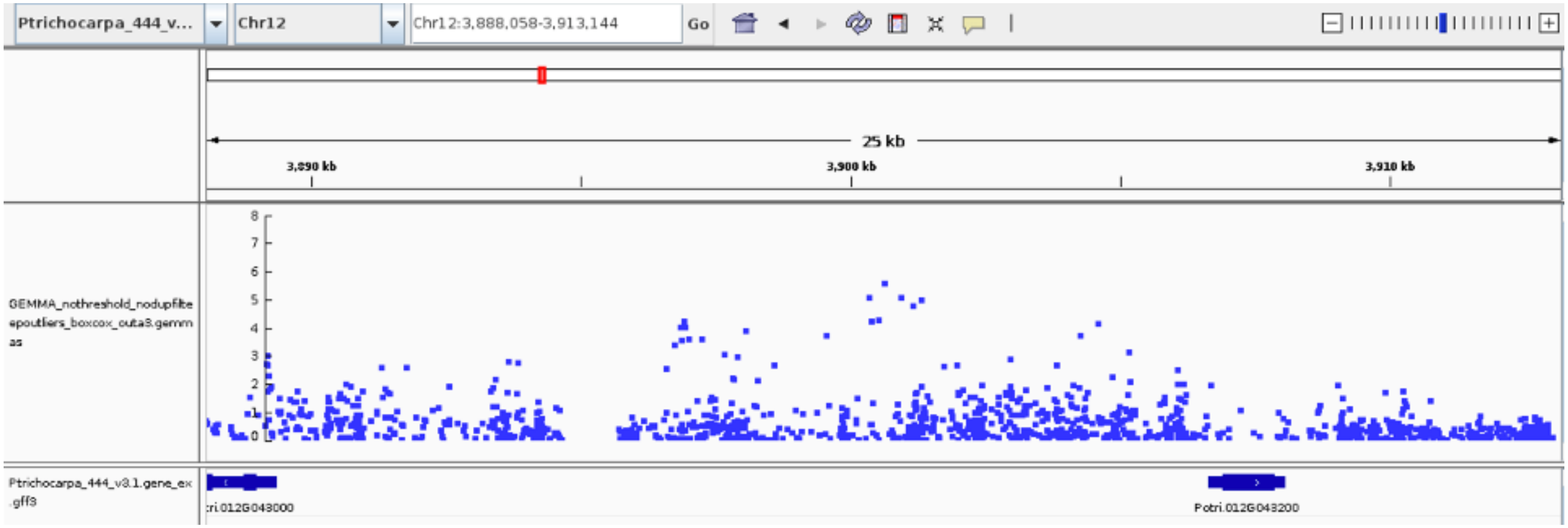
Ptrichocarpa_444_v...
Chr12
Chr12:3,888,058-3,913,144
Go
25 kb
3,890 kb
3,900 kb
3,910 kb
GEMMA_nothreshold_nodupfiltepoutliers_boxcox_outa3.gemmas
Ptrichocarpa_444_v3.1.gene_ex.gff3
:ri.012G043000
Potri.012G043200

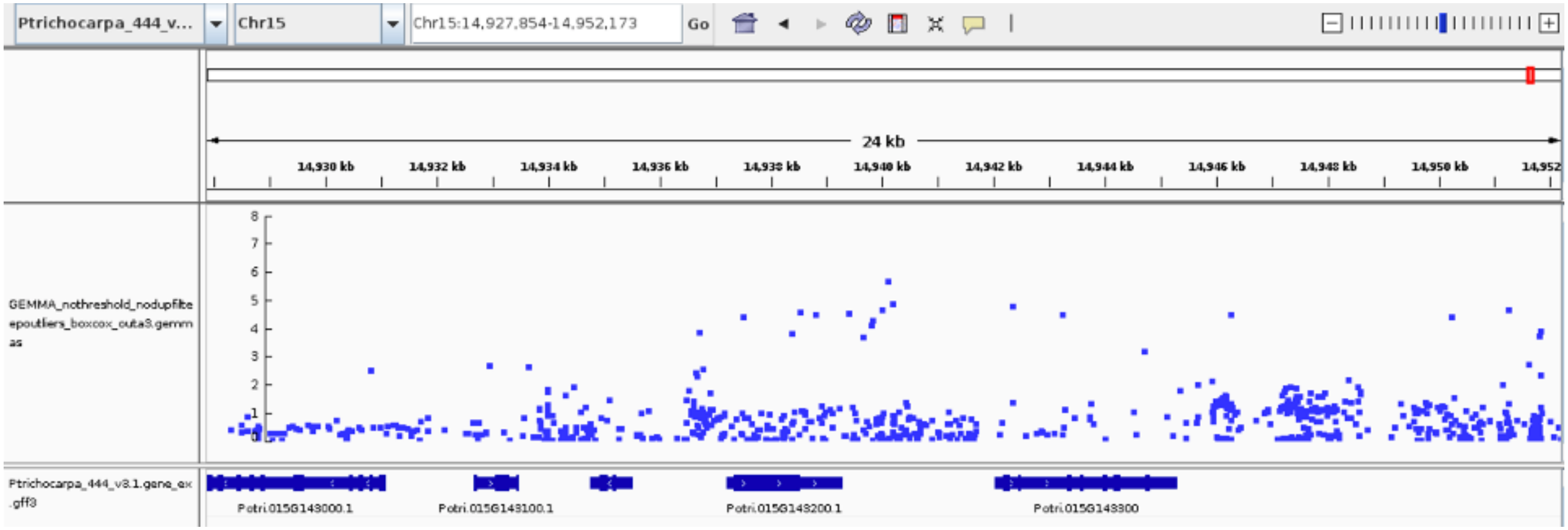
Ptrichocarpa_444_v...
Chr15
Chr15:14,927,854-14,952,173
Go
24 kb
14,930 kb
14,932 kb
14,934 kb
14,936 kb
14,938 kb
14,940 kb
14,942 kb
14,944 kb
14,946 kb
14,948 kb
14,950 kb
14,952
GEMMA_nothreshold_nodupfiltepoutliers_boxcox_outa3.gemmas
Ptrichocarpa_444_v3.1.gene_ex.gff3
Potri.015G143000.1
Potri.015G143100.1
Potri.015G143200.1
Potri.015G143300

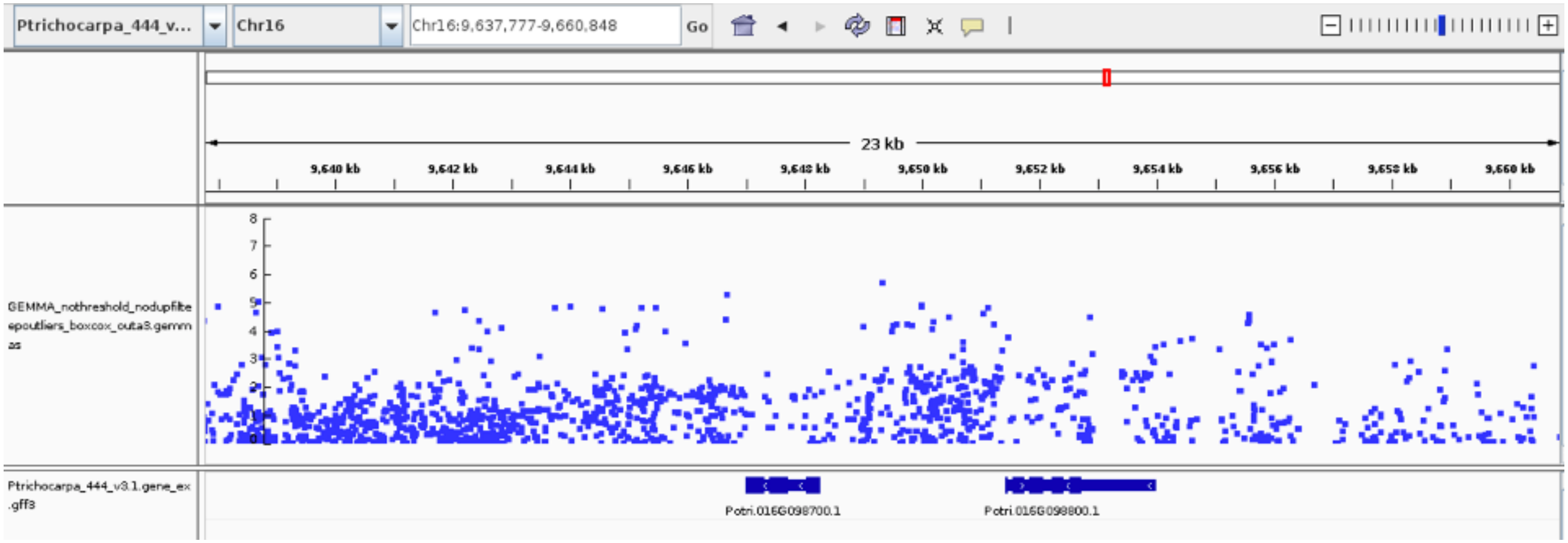
Ptrichocarpa_444_v...
Chr16
Chr16:9,637,777-9,660,848
Go
23 kb
9,640 kb
9,642 kb
9,644 kb
9,646 kb
9,648 kb
9,650 kb
9,652 kb
9,654 kb
9,656 kb
9,658 kb
9,660 kb
GEMMA_nothreshold_nodupfiltepoutliers_boxcox_outa3.gemmas
Ptrichocarpa_444_v3.1.gene_ex.gff3
Potri.016G098700.1
Potri.016G098800.1
