## Supplemental Methods; Supplemental Results and Discussion for "GWAS identifies candidate regulators of in planta regeneration in Populus trichocarpa"

Supplementary methods and materials

*Association Mapping*

Prior to GEMMA, SNPs were filtered based on minor allele frequency (MAF) > 0.01 and a missing rate of given SNPs across genotypes > 0.10 using PLINK (Purcell *et al.*, 2007), resulting in ~13.2 million SNPs. Filtering criteria were based on the whole SNP set of 1323 genotypes. GEMMA was used to compute Wald *p*-values for SNP effects, using the `-lmm 1` option. The number of SNPs analyzed by GEMMA was further reduced to ~12.5 million because GEMMA performed additional filtering with a default $R^2$ threshold of 0.9999 and default MAF and missing rates of 0.01 and 0.10, which were recomputed at this stage. In addition to performing association mapping, GEMMA was used to provide an estimate of narrow-sense SNP heritability ($h^2_{SNP}$) for each trait. Downstream GWAS and gene candidate evaluation was performed for traits with estimated $h^2_{SNP}$ above 0.10.

Second, we utilized logistics models for binarized traits with Generalized Mixed Model Association Test (GMMAT; (Chen *et al.*, 2016). Due to the computational expense of computing Wald *p*-values via logistic regression, we first performed the GMMAT variance component score test (`glmm.score`) for a genome-wide screen and then extracted a subset of 100 or 1,000 SNPs with the lowest score test *p*-values from each run and computed Wald *p*-values for these (using `glmm.wald`). This GMMAT workflow was performed with two SNP subsets prepared by PLINK: one had a missing rate threshold of 0.10 and an MAF threshold of 0.05 (~7.7 million SNPs), and the second had the same missing rate threshold but an MAF threshold of 0.01 (~13.2 million SNPs). Since rare SNPs led to inflated score tests and interfered with the ability to robustly select SNPs for downstream Wald tests, the former SNP set was relied upon to study six traits: wk. 2 callus area, wk. 3 callus area, wk. 2 shoot area, shoot PC1, shoot PC2 and callus/shoot PC1.

Finally, for multiple-marker tests we applied the SNP-set (Sequence) Kernel Association Test (SKAT; Ionita-Laza *et al.*, 2013) with untransformed traits. SKAT was performed on overlapping 3kb windows staggered by 1kb, using a set of 34.0 M SNPs filtered for a missing rate of 15%. The R extension Multi-Threaded Monte Carlo SKAT (MTMCSKAT) was used to run SKAT on a high-performance cluster, COMET (made available through NSF XSEDE

(Towns *et al.*, 2014). We calculated empirical *p*-values for top associations to avoid Type I and Type II error resulting from the non-normal distributions of untransformed traits. Two means of controlling for population structure were tested and compared with this workflow. We compared a "P" model in which structure is represented by principal components derived from SNPs (computed with PLINK) to a "Q" model in which structure is alternatively represented by subpopulation estimates produced by fastSTRUCTURE (Raj *et al.*, 2014).

To produce PCs for the P model, we employed a filtered set of ~10.3M SNPs with MAF > 0.05 and consulted scree plots and used K-means clustering to inform about the number of PCs appropriate for representing population structure; as a result we used 6 PCs for the P model.

To produce a Q matrix for use with SKAT Q models, we used fastSTRUCTURE using a subset of ~72k SNPs filtered based on LD, MAF, and missing rate using PLINK with parameters ` --indep-pairwise 100kb 10 0.05 --maf 0.05 --geno 0.1`. Ten replicates were performed with fastSTRUCTURE for each possible number of subpopulations (K) ranging from 3 to 12. To understand subpopulations in an evolutionary context, we used SNPhylo (Lee *et al.*, 2014) to produce a dendrogram from our SNP data. SNPhylo was run with a subset of ~129k SNPs prepared by PLINK with parameters ` --indep-pairwise 10kb 10 0.05 --maf 0.05 --geno 0.1`.

Geographical locations (longitude and latitude) were recorded for 1,301 of 1,323 genotypes in the SNP set and plotted against traits, SNP-derived PCs (for SKAT "P" model), primary subpopulation information (for SKAT "Q" model), and dendrogram information (from SNPhylo) using the `phylo.to.map` function in Phytools (R) and Google Maps "My Maps". Phytools was also used to cross-reference dendrograms with traits, SNP-derived PCs and primary subpopulation information (using function `phylo.heatmap`; Revell, 2012).

To inform about the appropriate window size for SKAT, as well as to inform about the likelihood of genes proximal to associated SNPs or SNP windows being directly involved in affecting traits (vs. being associated as a result of genetic linkage), we evaluated LD decay. To facilitate efficient computation of LD decay, a reduced SNP set (~78k SNPs) was prepared by PLINK with parameters `--maf 0.05 –geno 0.1 –thin 0.01`. Further reduced SNP files were prepared with PLINK to only include genotypes in the "Oregon" and "California" subpopulations (named based on general location of most genotypes in each). PLINK was further used to compute pairwise LD between all SNPs on each given chromosome with each SNP set. Using R, the average LD for each possible distance (e.g. 1bp, 2bp, 3bp… up to 50kb) was computed and plotted for the whole population as well as each of the two selected subpopulations.

*Evaluation of relationships between regeneration traits, subpopulations and geography*

Following the identification of subpopulation structure when fastSTRUCTURE was used to produce covariates for the SKAT "Q" model, we aimed to further investigate the relationships between traits, geography and theoretical ancestral subpopulations to gain insights into the possible adaptive evolution of these regeneration traits. To this end, we used `lm` in R to construct linear models regressing each trait over latitude and the Q matrix featuring estimates of each theoretical ancestral subpopulation's contribution to each individual's genome (from fastSTRUCTURE). We then visualized relationships, latitude and subpopulation using `ggplot2` in R.

### Supplementary results and discussion

*Distinct ancestral subpopulations supported by population structure and phylogeography analysis*

Relationships between evolutionary clades, geography, and population structure suggest that *P. trichocarpa*, despite its dioecy and long-distance gene flow, exists with a number of subpopulations that are statistically distinct albeit highly admixed. A total of 120 fastSTRUCTURE runs were performed, including 10 replicates for each value of K (subpopulation number) ranging from 2-13. The log marginal likelihood appears to be maximized with K equal to 6 or 7 (Fig. S5). For each individual in the population, the most closely related subpopulation was extracted and considered the primary subpopulation. Geographic and evolutionary patterns were revealed by cross-referencing of a dendrogram (SNPhylo; Lee *et al.*, 2014) with primary subpopulation and geographic location. These plots were evaluated with primary subpopulations from fastSTRUCTURE models both with K=6 and K=7 (Fig. S6); the K=7 model showed the strongest alignment between phylogeny and geography. Approximately from Seattle northward, individuals display a heavy degree of admixture and fail to cluster into clear subpopulations. Otherwise, the existence of several subpopulations is supported by agreement between phylogenetic clades, geographic location, and primary subpopulation label from fastSTRUCTURE. These include distinct subpopulations in the western region of Idaho and nearby eastern Oregon and Washington (and extending all the way to the eastern Washington Cascades near Yakima), the Willamette Valley of central western Oregon and nearby Western Washington, southwest Oregon and nearby northern California, northwestern Washington extending into southwestern Canada, and central western to northwestern Canada (Fig. S7). Also of note, we found evidence that LD rates vary across groups of distinct theoretical ancestral subpopulations, as shown by LD curves fit for "Oregon" and "California" groups (named by approximate location of primary theoretical ancestral subpopulation; Fig. S4). A linear model fit over this data for both groups, with an interaction term between primary ancestral subpopulation and a spline function of LD decay, indicated that this difference was statistically significant (empirical $p$-value < 0.001, 1000 permutations).

We further attempted to summarize population structure by performing PCA over SNP data using PLINK. Similar to fastSTRUCTURE subpopulation estimates, PCs explaining a substantial portion of variance show clear relationships with geography and most of the same phylogenetic clades (Fig. S8, S9). The use of six PCs to represent population structure in SKAT models, as discussed below, was supported by the scree plot (Fig. S10) and the relatively minor contributions of subsequent PCs to k-means clusters computed from PCs (Fig. S11).

We attempted to gain insights into the possible role of evolution in regeneration traits via relationships between the traits, latitude and theoretical ancestral subpopulation (Methods). At $\alpha = 0.005$, there appears to be a significant effect of latitude of clone origin on the trait of callus area at week four, while controlling for subpopulation. Several other relationships are significant at 0.05, between various callus traits and latitude and/or subpopulation (Table S3). Visualization of the relationships between traits and latitude along with regression trendlines showed a positive relationship between many regeneration traits and increasing latitude, but the significance of these trends was lost when considering clones of each given primary subpopulation independently (Fig. S12). Considering the lack of independence between variables of theoretical ancestral subpopulation and latitude, we advise caution in overinterpreting these results as evidence of either genetic draft or an adaptive role of regeneration, but also note several significant or borderline-significant trends indicating such a role may exist.

*Distinct subpopulations correlate with phylogeography*

The existence of distinct ancestral subpopulations of *P. trichocarpa* and a relationship of these subpopulations with geography is supported by cross-referencing of results from population structure analysis (fastSTRUCTURE), phylogenetics (SNPhylo), and geographical information for genotypes. These distinct subpopulations appear clearly in the southern portion of the population, whereas the northern portion displays a remarkable degree of admixture with mixed origins across the southern subpopulations. We speculate that, following the establishment of distinct southern subpopulations during the Last Glacial Period (Armstrong *et al.*, 1965), the recession of glaciers allowed for these subpopulations to spread to the northern region—where there has not yet been sufficient time or subdivision for distinctive populations to form. In contrast, the disjunct nature of many of the southern population groups is likely to have provided historical opportunities for differentiation. While previous work using approximately 12 isozyme loci did not reveal distinct subpopulations of *P. trichocarpa* over a more narrow, but similar geographical range (Weber & Stettler, 1981), our work demonstrates the much-increased power of genome-scale SNP data—where millions of loci are surveyed—to detect subpopulations.

*Similar results from SKAT with either PC or fastSTRUCTURE covariates*

We compared results from complementary SKAT models with population structure represented either by the fastSTRUCTURE Q matrix with 7 subpopulations (“Q model”) or by the first 6 PCs (“P model”) for a subset of four traits (callus area at wk. 4 and wk. 5; shoot area at wk. 4 and wk. 5). These models displayed a remarkable level of agreement, especially for *p*-values that met thresholds of significance and were thus selected for validation by computing empirical *p*-values with MTMCSKAT (Fig. S13).

*Overlap with genes implicated from published GWAS analyses of regeneration*

The candidates we identified showed very little similarity to results from related work. In prior work, GWAS was performed in 280 genotypes of *P. trichocarpa* to study traits related to in vitro callus regeneration. This study yielded eight candidate genes, none of which appear among our results (Tuskan *et al.*, 2018). A GWAS of traits related to roots and vegetative shoots in *Populus deltoides* × *simonii* with 434 genotypes produced 233 QTLs and multiple gene candidates were considered within proximity of each QTL, yielding a total of 595 unique gene candidates, only three of which were also found among traits analyzed in our study. Potri.015G018200, encoding a putative protein kinase, is a gene candidate from our analysis of callus area at week two as well as a prior analysis of a measurement of the number of leaves per vegetative shoot in *P. euphratica*. This leaf number trait also yields an association for Potri.004G156900, encoding a putative RETICULATA-related protein also appearing as a candidate in our analysis of shoot area at week four. Another association is with Potri.019G035200, which encodes an oxygenase involved in heme degradation within chloroplasts; it was found among our gene candidates for callus at week two as well as in the same work for average stem diameter (Sun *et al.*, 2019).

In a review of GWAS of regeneration in diverse species, Lardon and Geelen (Lardon & Geelen, 2020) noted that gene candidates identified across studies are non-overlapping to a great extent. Some of the potential causes for the low degree of overlap include genetic differences between study populations, variation in tissue or explant physiology, variation in the treatments used to promote regeneration, random variation in detection given underpowered statistics and numerous genes under polygenic traits control, and differing statistical approaches (Lardon & Geelen, 2020). All of these factors would apply to our study vs. the other published work in *Populus*. Another likely contributor to lack of overlap is that our GWAS is the only one studying in planta regeneration, as opposed to in vitro regeneration or vegetative shoot development, and the genetic control of these developmental processes is likely to vary significantly. Finally, we note that the traits obtained from our computer vision pipeline are distinct from those in these prior studies, which made use of various manual scoring systems.
